# Voltage-dependent anion channels are mitophagy receptors mediating the recycling of depolarized mitochondria in *Arabidopsis*

**DOI:** 10.1101/2025.05.29.656919

**Authors:** Wenlong Ma, Juncai Ma, Kaiyan Zhang, Xuanang Zheng, Pengfei Wang, Lanlan Feng, Shizhong Ming, Xiaohong Zhuang, Jun Zhou, Caiji Gao, Byung-Ho Kang

## Abstract

The mitochondrion is an essential organelle in eukaryotic cells, playing crucial roles in cellular respiration and intracellular signaling pathways. To maintain a healthy population of mitochondria, dysfunctional and excess mitochondria are selectively removed through an autophagic process known as mitophagy. Over the past few decades, various autophagy-related (ATG) proteins involved in mitophagy have been well characterized in yeast and mammalian cells since it has significance to the survival of eukaryotes. While the core autophagy machinery responsible for autophagosome formation is conserved among eukaryotes, the homologs of key regulators of mammalian system is absent in plant. In this study, we identified a unique mitophagy mechanism in plant, that three voltage-dependent anion channel (VDAC) family proteins in the mitochondria outer membrane — specifically VDAC1, VDAC2, and VDAC3 — as mitophagy receptors in *Arabidopsis*. These proteins were required for translocation of ATG8 from the cytosol to the mitochondria surface, when *Arabidopsis* cells were treated with an uncoupler, 2,4-dinitrophenol (DNP). The VDACs interacted directly with ATG8 through an ATG8-interacting motif (AIM) located in their amino (N) termini. Furthermore, *vdac* mutants exhibited impaired uncoupler-induced mitophagy and accumulated damaged mitochondria. These mitophagy-related phenotypes were more pronounced in *vdac* double and triple mutant lines. Altogether, our results indicated that VDAC1, 2, and 3 recruit ATG8 to depolarized mitochondria, facilitating the formation of mitophagosomes, presenting a distinguishing mitophagy pathway with mammalian system.

## Introduction

Autophagy contributes to the maintenance of cellular homeostasis by recycling dysfunctional or surplus cellular components (Lamark and Johansen, 2021). Macroautophagy (referred as autophagy hereafter) is a type of autophagy involving the formation of double membrane structures known as autophagosomes. These autophagosomes sequester the cytosol, organelles, protein aggregates, or pathogens in their lumen and subsequently fuse with vacuoles/lysosomes to deliver their contents for degradation (Ohsumi, 2014). The process of autophagy is orchestrated by a set of core machinery termed autophagy-related (ATG) proteins, which are evolutionarily conserved across species (Lamark and Johansen, 2021). Among those ATGs, ATG8 has been monitored as a marker for autophagosome formation because ATG8 undergoes cleavage and lipidation for its recruitment to autophagosomes and stays with the double membrane till lysosomal/vacuolar degradation (Marshall et al., 2019).

Autophagy operates in either selective or non-selective modes, depending on its substrate selectivity. Non-selective autophagy indiscriminately engulfs cytoplasmic components for bulk degradation (Su et al., 2020). Selective autophagy relies on the recognition of receptors that bind directly to their cargos or ubiquitin modifications on cargos (Otegui et al., 2024). These receptors act as molecular adaptors, linking the cargo to the autophagic machinery, often through interactions with ATG8. Autophagy receptors possess short amino acid stretches called ATG8-interacting motif (AIM) or LC3-interacting region (LIR), mediating the binding of receptors to ATG8 that facilitates the selective engulfment of substrates into autophagosomes.

Damaged or excess mitochondria are selectively recycled by mitophagy for keeping the mitochondrial population healthy (Palikaras et al., 2018). The PINK1/Parkin-dependent pathway is arguably the most extensively studied mitophagy process in mammalian cells to eliminate damaged mitochondria (Harper et al., 2018). PINK1 is a protein kinase recruits and phosphorylates an E3 ubiquitin ligase, Parkin, on the surface of damaged mitochondria (Gan et al., 2022). Proteins ubiquitinated by Parkin then attract mitophagy receptors such as p62, OPTN, and NDP52, which link the ubiquitinated mitochondrial proteins with ATG8 (Harper et al., 2018). Certain outer mitochondrial membrane (OMM) proteins like BNIP3, NIX, and FUNDC1 directly interact with ATG8, promoting mitophagosome formation upon the loss of mitochondrial membrane potential (Liu et al., 2012). Furthermore, Prohibitin 2 (PHB2), an inner mitochondrial membrane (IMM) protein, can act as a receptor interacting with ATG8 when the OMM has been ruptured (Wei et al., 2017). Moreover, cardiolipin, an IMM-located phospholipid, can translocate to the OMM to recruit ATG8 (Chu et al., 2013). Lastly, mtDNA released by damaged mitochondria can be removed by t mitochondrial transcription factor A (TFAM) (Liu et al., 2024).

Although mitophagy in plant cells has not been investigated so much as in animal cells, it has been demonstrated that mitophagy operates to degrade depolarized mitochondria in plants, which is critical for plant growth and survival (Broda et al., 2018). *Arabidopsis* root cells challenged with an uncoupler, 2,4-dinitrophenol (DNP), and cotyledon cells during de-etiolation accumulate depolarized mitochondria, which are subsequently eliminated by mitophagy (Ma et al., 2021; Wijerathna-Yapa et al., 2021). An *Arabidopsis* mitochondrial clustered family protein, friendly mitochondria (FMT), is involved in the mitophagosome formation and its absence delays chloroplast biogenesis during de-etiolation (El Zawily et al., 2014; Ayabe et al., 2021; Ma et al., 2021). In addition, ATG11 was shown to be implicated in mitophagy in senescing *Arabidopsis* leaf cells (Li et al., 2014). Outer mitochondria membrane proteins, TRB1 and 2, bring ATG8 and the endoplasmic reticulum (ER) elements together to facilitate mitophagosome formation (Li et al., 2022). Furthermore, mitochondria-localized FCS-like zinc finger (FLZ) proteins interact with ATG8 during carbon starvation in plants (Yang et al., 2023). However, our understanding as to the molecular factors playing roles in the selective mitochondria degradation in plant cells remains limited.

Voltage-Dependent Anion Channels (VDACs) are β-barrel proteins embedded in the outer mitochondrial membrane, that facilitate transport of anions between the cytosol and mitochondria (Messina et al., 2012; Di Rosa et al., 2021). VDACs play roles in cellular stress responding processes, including mitophagy and apoptosis (Shoshan-Barmatz et al., 2008; Kusano et al., 2009; Shoshan-Barmatz et al., 2010). Human VDAC1 is poly-ubiquitinated at its lys27 by Parkin and serves as a surface mark for recruiting p62 to mitochondria, initiating the formation of mitophagosome (Geisler et al., 2010). Interestingly, a recent study reported that the leafhopper VDAC1 can directly interact with ATG8 as an OMM receptor of mitophagy (Chen et al., 2023). On the other hand, mono-ubiquitin modification to VDACs acts as a signal for apoptosis (Ham et al., 2020). Oligomerization and conformational changes of VDACs are linked to stressed mitochondria (Abu-Hamad et al., 2009; Shoshan-Barmatz et al., 2010; Kim et al., 2019; Ren et al., 2023). Altogether, these lines of evidence suggest that VDACs have functions in multiple aspects in stress responses.

Here, we report *Arabidopsis* VDAC1, 2, and 3 as mitophagy receptors recruiting ATG8 to depolarized mitochondria. In a proteomic analysis of *Arabidopsis* cells expressing yellow fluorescent protein-tagged ATG8e (YFP-ATG8e), VDAC1, 2, and 3 were pulled down after the plants were incubated with DNP. The VDACs have flexible N-terminal domains involved in the regulation of channel permeability in response to changes in mitochondrial membrane potential. Our data show that the three VDACs have a conserved AIM in their N-terminus critical for recruiting ATG8 to depolarized mitochondria and that inactivation of genes encoding the VDACs led to accumulation of aberrant mitochondria in DNP treated *Arabidopsis* cells.

## Results

### VDAC isoforms are involved in DNP-induced mitophagy in *Arabidopsis* root cells

To identify proteins associated with ATG8 during mitophagy, we conducted an immuno<u>p</u>recipitation-<u>m</u>ass <u>s</u>pectrometry (IP-MS) screen using a transgenic *Arabidopsis* line expressing YFP-ATG8e. Seedlings expressing YFP-ATG8e were subjected to DNP treatment, and proteins interacting with YFP-ATG8 were isolated using GFP-affinity beads (Figure 1A and 1B). Control experiments included seedlings expressing free YFP or YFP-ATG8e with DMSO treatment. Volcano plots revealed that 510 proteins were significantly enriched in DNP-treated YFP-ATG8e compared to the YFP control, and 606 proteins were enriched in DNP-treated YFP-ATG8e in comparison with the DMSO-treated mock group, with an overlap of 275 enriched proteins from the two groups (Figure 1C). Notably, some of the core ATG proteins such as ATG3 and ATG7 were among the overlapping group. Gene Ontology analysis indicated that mitochondria, chloroplasts, and Golgi stacks were among the top three membrane-bound organelles where the 275 proteins are localized (Figure 1D). Considering their enrichment in IP-MS results regarding to OMM localization, we identified VDAC1, 2, and 3 as potential candidates for mitophagy key factors that may recruit ATG8 to mitochondria.

**Figure 1.**
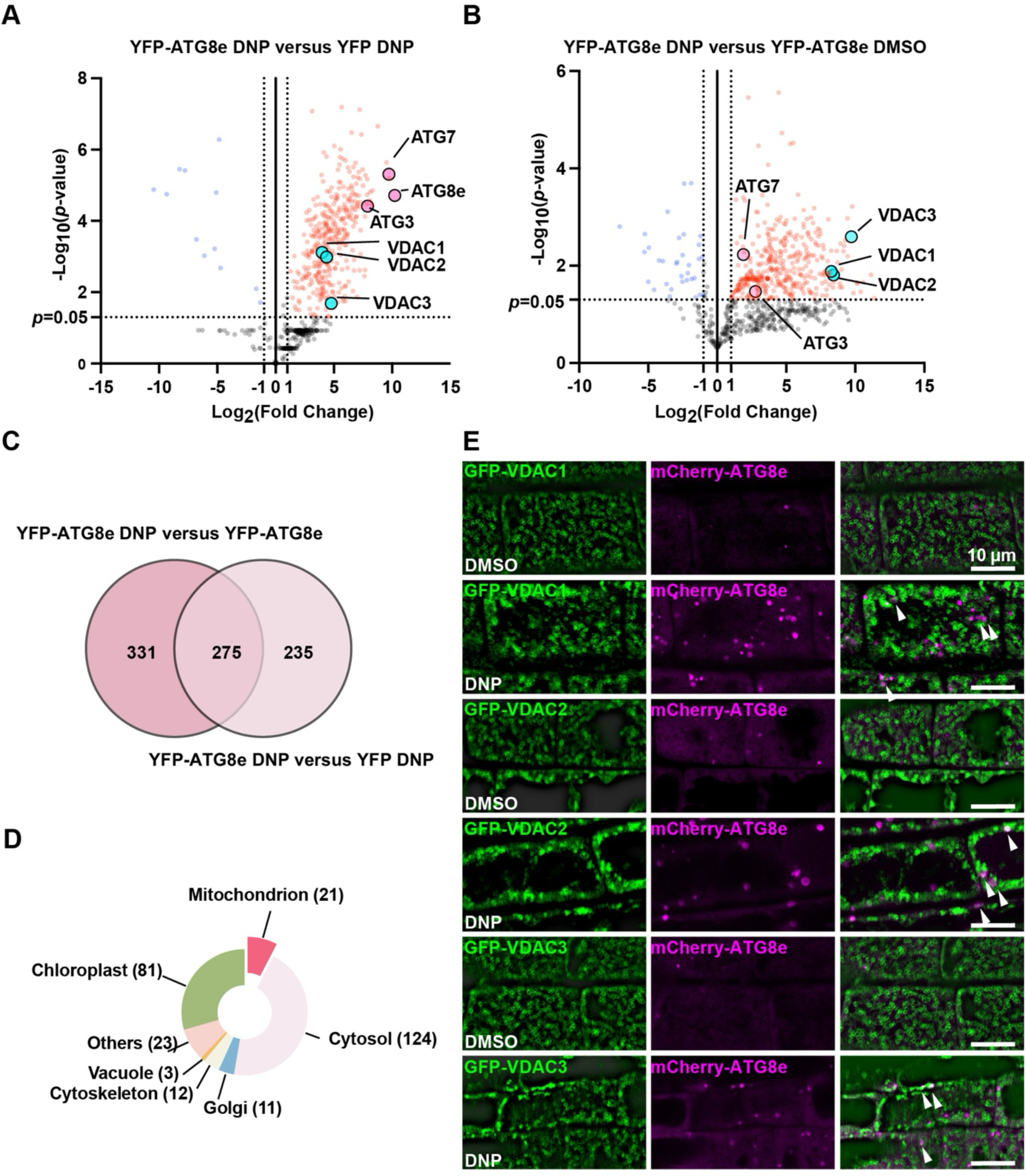
VDAC1, 2, and 3 are associated with autophagosomes after depolarization stress. A-B) Volcano plots showing the co-immunoprecipitation proteomics comparing DNP-treated YFP-ATG8e with DNP-treated YFP by **A)**, and DNP-treated YFP-ATG8e with DMSO-treated YFP-ATG8e by **B)**. X- and Y-axis display log_2_(fold change) and −log_10_(*p*-value), respectively. Dashed lines represent threshold for log_2_(fold change) = ±1 and *p*-value = 0.05. VDAC1, VDAC2, and VDAC3 identified in the comparison between DMSO- and DNP-treatment are marked with pink dots. Core autophagic proteins are denoted with cyan dots. n=3. **C)** Venn diagram illustrating significantly enriched proteins in both of the two proteomic screening in **A)** and **B)**. **D)** Pie chart showing the subcellular locations of the 275 proteins in the overlapped group of the Venn diagram in **C)**. Subcellular localization of the proteins were assigned with the Gene Ontology tool. Numbers of proteins for each organelle are indicated in the parentheses. **E)** Confocal micrographs of *Arabidopsis* root cells expressing GFP-VDAC and mCherry-ATG8e after incubation with DMSO or DNP. White arrowheads mark mitochondria (VDACs) associated with mCherry-ATG8e. Scale bars: 10 µm.

To confirm the association of VDAC1, 2, and 3 with ATG8, we generated transgenic *Arabidopsis* lines expressing <u>g</u>reen fluorescent <u>p</u>rotein (GFP) fused to the N-terminus of VDAC1, 2, or 3 (GFP-VDAC) and crossed the lines to the line expressing mCherry-tagged ATG8e. The GFP-VDACs were localized to the mitochondrial surface and associated with mCherry-ATG8e puncta when the transgenic seedlings were incubated with DNP (Figure 1E). These findings are consistent with the association of ATG8 with VDAC1, 2, or 3 triggered by membrane depolarization in the IP-MS dataset.

### VDAC1, 2, and 3 directly interact with ATG8s as autophagic receptors

To test the potential roles of VDAC1, 2, and 3 as mitophagic receptors, we examined whether they bind to ATG8 directly. Co-overexpression of hexa-histidine-fused VDACs (6×His-VDACs) and mCherry-ATG8e in *Arabidopsis* seedlings showed positive binding in *in vivo* co-IP assay using Ni-NTA beads in a DNP dependent manner (Figure 2A). To verify the IP result, we carried out *in vitro* pull-down assay in which recombinant protein samples from *E. coli.* lines expressing Glutathione S-transferase-fused VDACs (GST-VDACs) or 6×His-tagged ATG8e (6×His-ATG8e) were purified and incubated together. 6×His-ATG8e were pulled down together with GST-VDAC1, 2, and 3 by GST-affinity beads, supporting that VDAC1, 2, and 3 directly interact with ATG8e (Figure 2B). When GFP-VDACs were co-expressed with mCherry-ATG8e in the *Arabidopsis* protoplasts, all three VDAC colocalized with mCherry-ATG8e, while free GFP failed to colocalised with ATG8 puncta (Figure 2C). The interactions were further confirmed by yeast two-hybrid (Y2H) assay. Each of the nine *Arabidopsis* ATG8 isotypes (AtATG8a-i) was individually tested with the three VDAC proteins. The pairwise tests demonstrated that VDAC1, 2, and 3 interacted with all nine AtATG8 proteins in the Y2H system (Figure 2D). These findings provide evidence that VDAC1, 2, and 3 directly interact with ATG8, supporting their potential role as OMM receptors for mitophagy.

**Figure 2.**
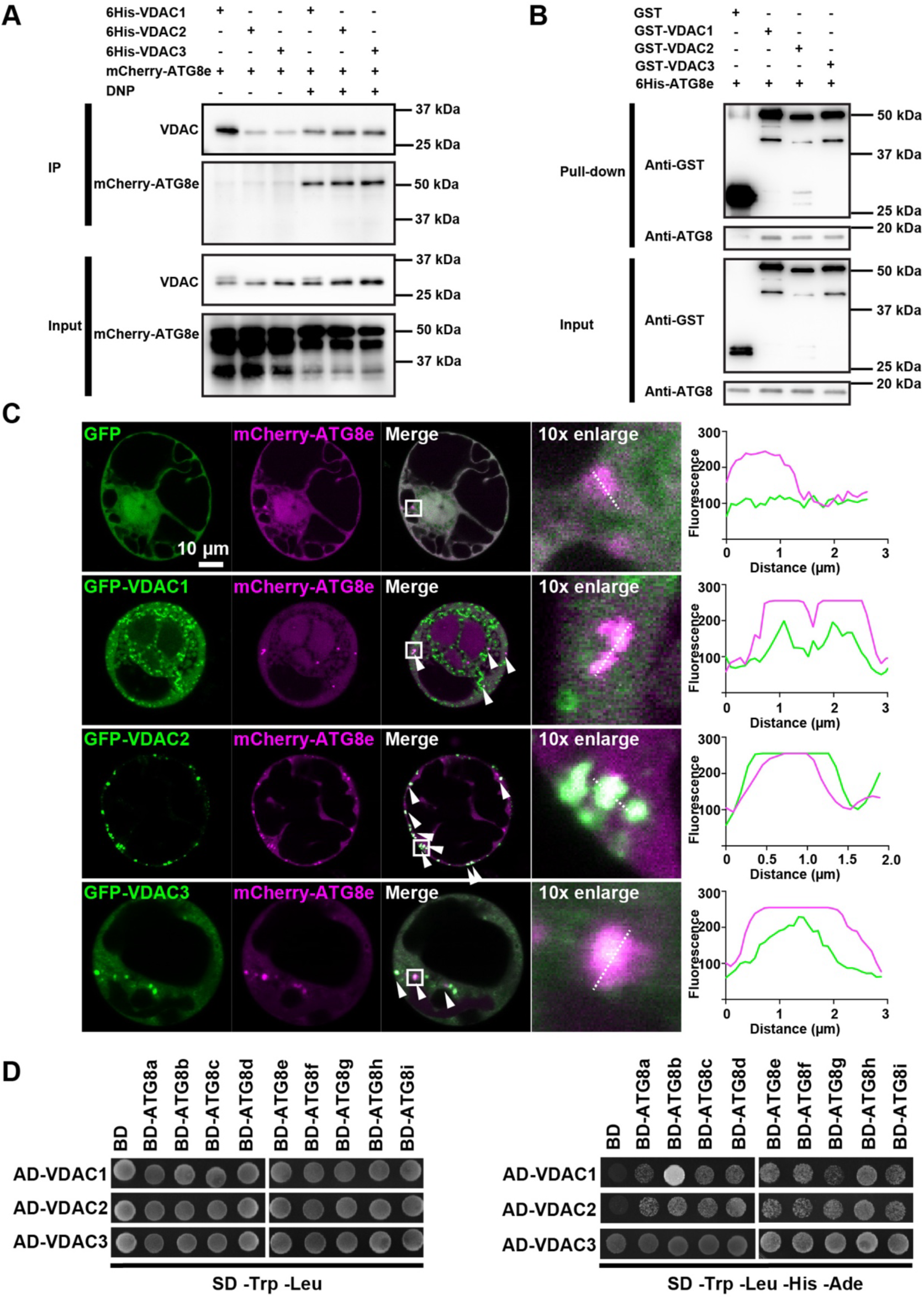
VDAC1, 2, and 3 directly interacts with ATG8 isoforms. **A)** Co-immunoprecipitation (Co-IP) of *Arabidopsis* seedlings co-expressing 6×His-VDACs and mCherry-ATG8e. Seedlings of the transgenic line were treated with DMSO or DNP and protein samples from the seedlings were incubated with Ni-NTA beads for isolating proteins associated with 6xHis-VDACs. Protein isolates from the beads were subjected to immunoblot analysis. **B)** Pull-down assay of 6×His-ATG8e by purified GST-fused VDAC1, 2, and 3. Free GST was employed as a negative control. **C)** Confocal microscopy images of *Arabidopsis* protoplasts co-expressing GFP-VDACs with mCherry-ATG8e. Puncta where GFP and mCherry fluorescence colocalize are readily discerned (arrowheads). Fluorescence intensity profiles of GFP and mCherry overlapped along the dashed lines through such puncta in the 10X magnified views. Scale bar: 10 µm. **D)** Yeast-two-hybrid assay showing that VDAC1, 2, and 3 interact with all the nine isoforms of *Arabidopsis* ATG8 in. BD: binding domain, AD: activating domain, SD -Trp -Leu: Synthetic dextrose minimal medium without tryptophan and leucine, SD -Trp -Leu -His -Ade: Synthetic dextrose minimal medium without tryptophan, leucine, histidine, and adenine.

### ATG8-interacting motifs in the VDAC N-terminal domains bind to the LIR docking site

To characterize the interactions between VDACs and ATG8, firstly we tested which type the interaction is. *In vitro* pull-down results present that GST-VDAC interact with ATG8e via AIM-LDS interaction. When the LDS (Y51, L52), but not UDS (I78 - I80), in ATG8e were mutated by replaced with alanine, the interaction was abolished (Figure 3A). To find out the exact AIM in VDACs, we analyzed the VDAC amino acid sequences using the online iLIR Autophagy Database (https://ilir.warwick.ac.uk/) that predicts AIMs. Among AIMs suggested by iLIR (Supplementary Figure 1A), one at the N-terminus stood out for two reasons: (1) it is conserved among all three VDAC1, 2, and 3 but not in 4 and 5, (2) it is situated in the N-terminal overhang accessible for interaction with ATG8. Other AIMs were found not conserved among the three VDAC proteins and obscured in the transmembrane domains of the β-barrel proteins, not accessible to cytosolic proteins like ATG8 (Figure 1B). We simulated the interactions between the VDAC N-terminal overhang (VDAC-N) and ATG8 with AlphaFold2-multimer, an addition to AlphaFold2 designed for predicting protein-protein interactions. The side chain of Tyr8 (or Phe8) and Ile11 in the AIM of VDAC-N were calculated to fit into the W-site and L-site of ATG8’s LDS, respectively (Figure 3B). Moreover, *in silico* mutation of the AIM’s key amino acids to alanine disrupted the interaction (Figure 3C). *In vitro* pull-down assays confirm that GST-VDAC-N interact with ATG8e. When the AIM residues or LDS in ATG8e were mutated by replaced with alanine, the interaction was abolished, in agreement with the notion that VDAC-N mediates the interaction between VDAC1, VDAC2, VDAC3 and ATG8s (Figure 3D). Transient expression in protoplast and Y2H also indicated that N-terminal AIMs are critical for VDAC1, 2, and 3’s binding to ATG8 isotypes or colocalization with mCherry-ATG8e, respectively (Figure 3E and 3F).

**Figure 3.**
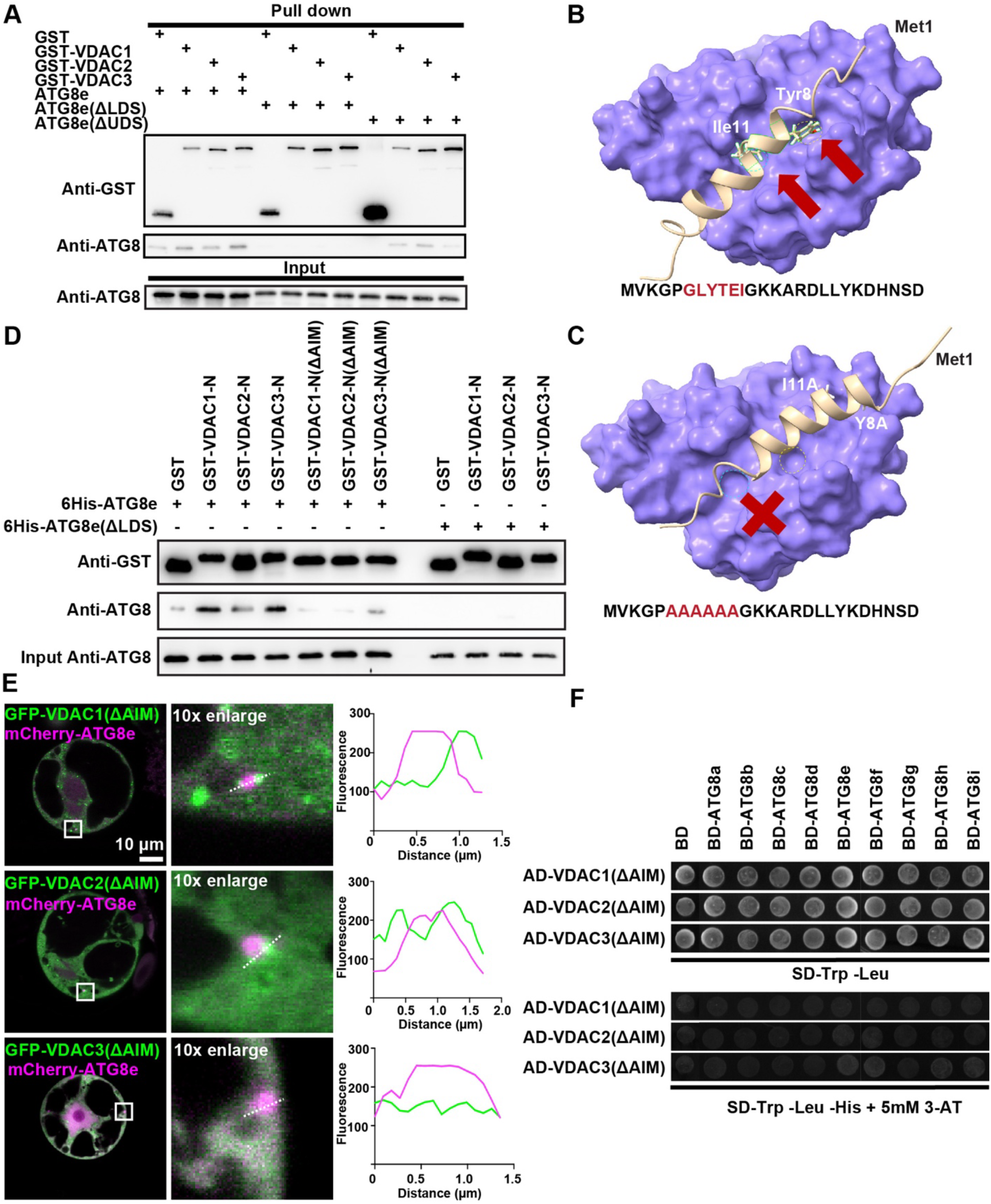
Binding of VDACs to ATG8 is mediated by via their conserved AIMs. **A)** Pull-down assay of between GST-fuse VDAC1, 2, 3 and ATG8e, ATG8e with LDS mutation (Y51A, L52A), or ATG8e with UDS mutation (I78A, F79A, I80A). Intact LDS domain of ATG8e is required for its binding with VDACs, while the mutation in the UDS domain did not affect the interaction. **B-C)** *In silico* simulation of ATG8e’s binding with the AIM in the VDAC1 N-terminal (VDAC1-N) domain by AlphaFold2-multimer. Side chains of Tyr8 and Ile11 in the VDAC-N domain fit in the W-site and L-site of the AIM (indicated with red arrows), respectively. When the six key amino acid residues of the AIM were replaced with alanine, ATG8e did not bind to mutated AIM of the VDAC1-N domain in AlphaFold2-multimer simulation **C)**. **D)** Pull-down assay indicating that AIMs in the VDAC-N domains interact with the LDS of ATG8e. Mutations in the AIM or LDS block the interaction. **E)** Confocal microscopy images of *Arabidopsis* protoplasts co-expressing GFP-VDACs without the AIM in their N-terminal domain (VDAC-ι1AIM) and mCherry-ATG8e. GFP and mCherry fluorescence did not colocalize. When fluorescence intensity profiles of closely localized GFP and mCherry puncta were compared, they failed to overlap, confirming that the AIMs are required for the VDACs-ATG8 interaction. Scale bar: 10 µm. **F)** Yeast-two-hybrid assay with VDAC constructs in which AIMs of the VDAC-N domain were mutated. BD: binding domain, AD: activating domain, SD -Trp -Leu: Synthetic dextrose minimal medium without tryptophan and leucine, SD -Trp -Leu -His: Synthetic dextrose minimal medium without tryptophan, leucine, histidine, 3-AT: 3-aminotriazole were added reducing background.

### VDAC N-terminal overhang became accessible to external proteases under mitochondria depolarization stress

It has been reported that the N-terminus of VDAC (VDAC-N) shows highly flexibility and plays a central role in channel gating according to alteration in mitochondrial membrane potential depolarization (Abu-Hamad et al., 2009; Shuvo et al., 2016). Therefore, we hypothesized that DNP treatment causes conformational changes in VDACs, making the transloction of VDAC-N and AIM in VDAC-N more accessible to ATG8 to facilitate mitophagosome assembly. To test the hypothesis, we designed a proteinase K digestion assay. If VDAC-N inside the transmembrane pore were to come out of the pore when mitochondria becomes depolarized, its digestion by proteinase K would be facilitated by DNP stress. As a positive control for proteinase K digestion, we adopted an OMM protein, TOM5, fused to a GFP in its domain facing the cytosol. As the GFP stays on the mitochondrial surface, it should readily be degraded by proteinase K. TOM9 served as a negative control since it is embedded from the intermembrane space side of OMM. Intensity of a mitochondria matrix protein, isocitrate <u>d</u>e<u>h</u>ydrogenase, IDH presents the amount of intact mitochondria. Lastly, Triton X-100 were added to solution to breakdown mitochondrial membrane to verify the proteinase K efficiency (Figure 4A).

**Figure 4.**
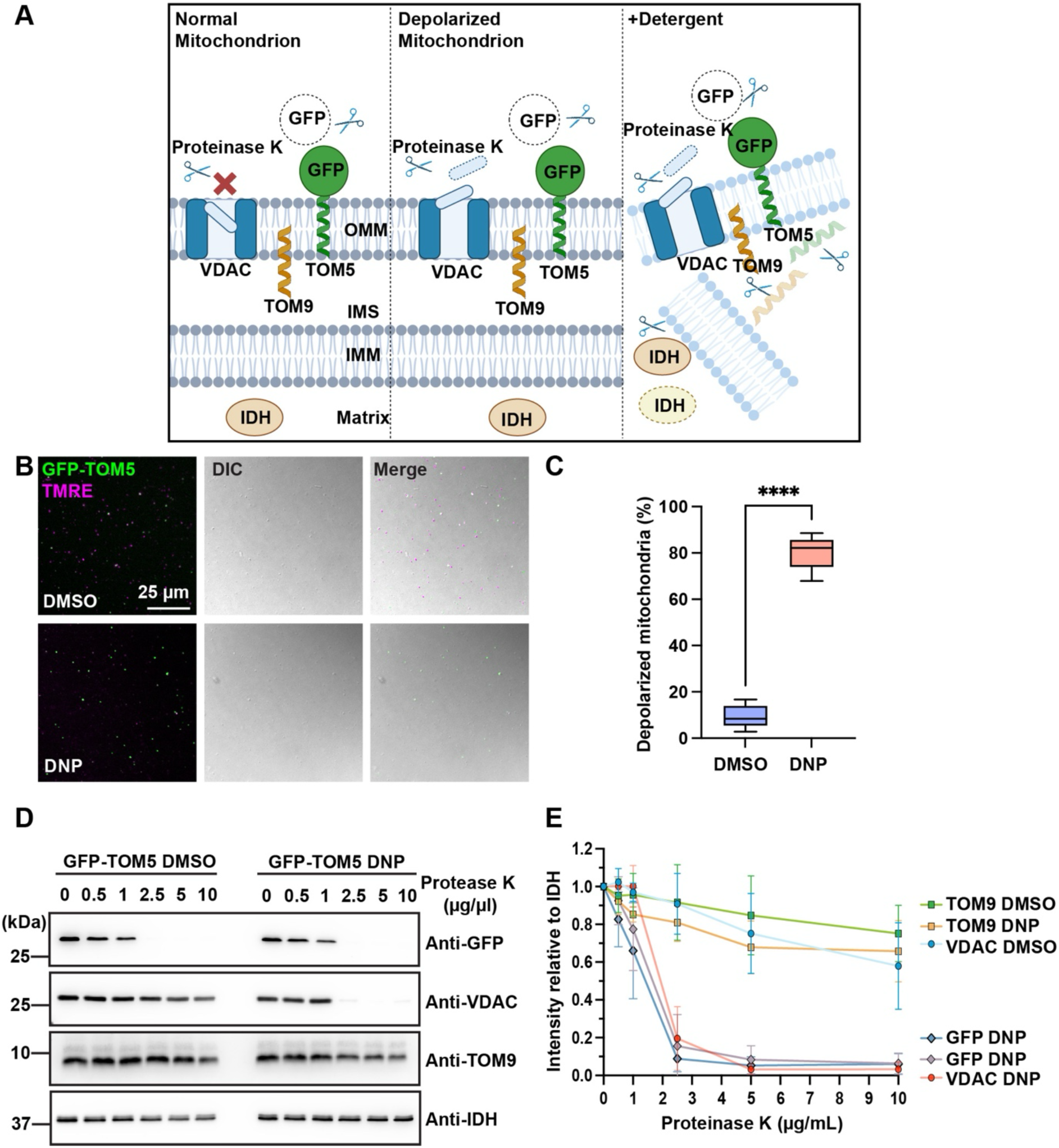
Depolarization of mitochondria facilitates degradation of VDACs’ N-terminal domain by protease K. **A)** A schematic diagram illustrating positions of the N-terminal domain of VDAC, TOM9, and GFP in the GFP-TOM5 fusion protein examined in our proteinase K digestion assay. VDACs were hypothesized to undergo a conformational change in depolarized mitochondria to expose their N-terminal domains to the cytosol, making them more susceptible to digestion by Proteinase K. TOM9 acts as a negative control as it faces the intermembrane space from the outer mitochondrial membrane (OMM). GFP-TOM5 serves as a positive control since its GFP is exposed to the cytosol. All protein amount is normalized based on the intensity of a mitochondrial matrix protein, isocitrate dehydrogenase (IDH). **B)** Confocal micrographs of mitochondria isolated from *Arabidopsis* seedlings expressing GFP-TOM5. Mitochondria were visualized with green fluorescence from GFP-TOM5. Note that majority of isolated mitochondria are positive for TMRE staining (DMSO panel) indicating that they retain their membrane potential. However, the TMRE fluorescence rapidly disappeared upon DNP treatment (DNP panel). DIC: differential interference contrast. Scale bar: 25 µm. **C)** Quantification of increases in depolarized mitochondria by DNP shown in **B)**. Means ± s.d.. n = 13 for each group. Unpaired t-test; *p* < 0.0001. **D)** Immunoblot analyses of mitochondrial proteins after proteinase K digestion. GFP in GFP-TOM5 and VDAC are digested quickly after DNP incubation. **E)** Quantification of proteinase K assay results from immunoblot analysis. Intensities of IDH were measured to estimate amounts of mitochondria in each experimental sample set. TOM9 amounts indicate the integrity of mitochondria outer membrane. GFP-TOM5 levels indicating the efficiency of proteinase K digestion. All protein amounts relative to IDH were normalized to the results from the 0 µg/ml Protease K sample. Means ± s.d.. n=3 for each group.

To processed *in vitro* digestion, 7-days-old seedlings expressing GFP-TOM5 were treated by DMSO or DNP. Purified mitochondria observed via confocal microscopy. GFP-TOM5 served as the mitochondria marker and DNP-induced depolarization was verified by tetramethyl rhodamine ethyl ester (TMRE), a fluorescent membrane potential sensitive dye for microscopy imaging. Mitochondria lacking membrane potential were enriched in DNP-treated samples more than sixfold than in DMSO-treated samples (Figure 4B and 4C). Our VDAC antibody recognized the N-terminal regions of all three VDACs but not their C-terminal regions, making it possible to monitor behaviors of the N-terminal domain with the antibody (Supplementary Figure 2A). When we incubated isolated mitochondria samples with serial dilutions of Protease K, degradation of VDACs’ N-terminus was significantly accelerated in DNP-treated mitochondria samples when compared with DMSO-treated samples (Figure 4C and 4E). Another OMM protein TOM9 did not exhibit such enhanced degradation under the same conditions. The levels of GFP, VDAC, and TOM9 were normalized with those of IDH. GFP, VDAC, TOM9, and IDH were completed eliminated when mitochondria membranes were disrupted with Triton X-100 (Supplementary Figure 2C).

To verify whether the exposure of VDAC N-terminal overhang in cytosol is sufficient for recruiting ATG8, we expressed artificial receptors in *Arabidopsis* protoplasts (Figure 5). The original or AIM-mutated versions (ΔAIM) of VDAC N-terminal (VDAC-N) were tagged with GFP and fused to TOM5. These chimeric OMM proteins were co-expressed with mCherry-ATG8e in *Arabidopsis* PSBD cells. In this architecture, VDAC N-terminal AIMs locate at the periphery of mitochondria. VDAC-Ns of VDAC1 (VDAC-1N), VDAC2 (VDAC-2N), or VDAC3 (VDAC-3N) recruited mCherry-ATG8e to mitochondria (Figure 5A and 5C), while VDAC-N(ΔAIM)- GFP-TOM5 did not (Figure 5B and 5C). The diagram explaining this construct is provided in Figure 5D. These results imply that ATG8 gains access to the N-terminal AIMs of VDAC1, 2, and 3 in depolarized mitochondria, promoting their recruitment therefore the assembly of mitophagosome.

**Figure 5.**
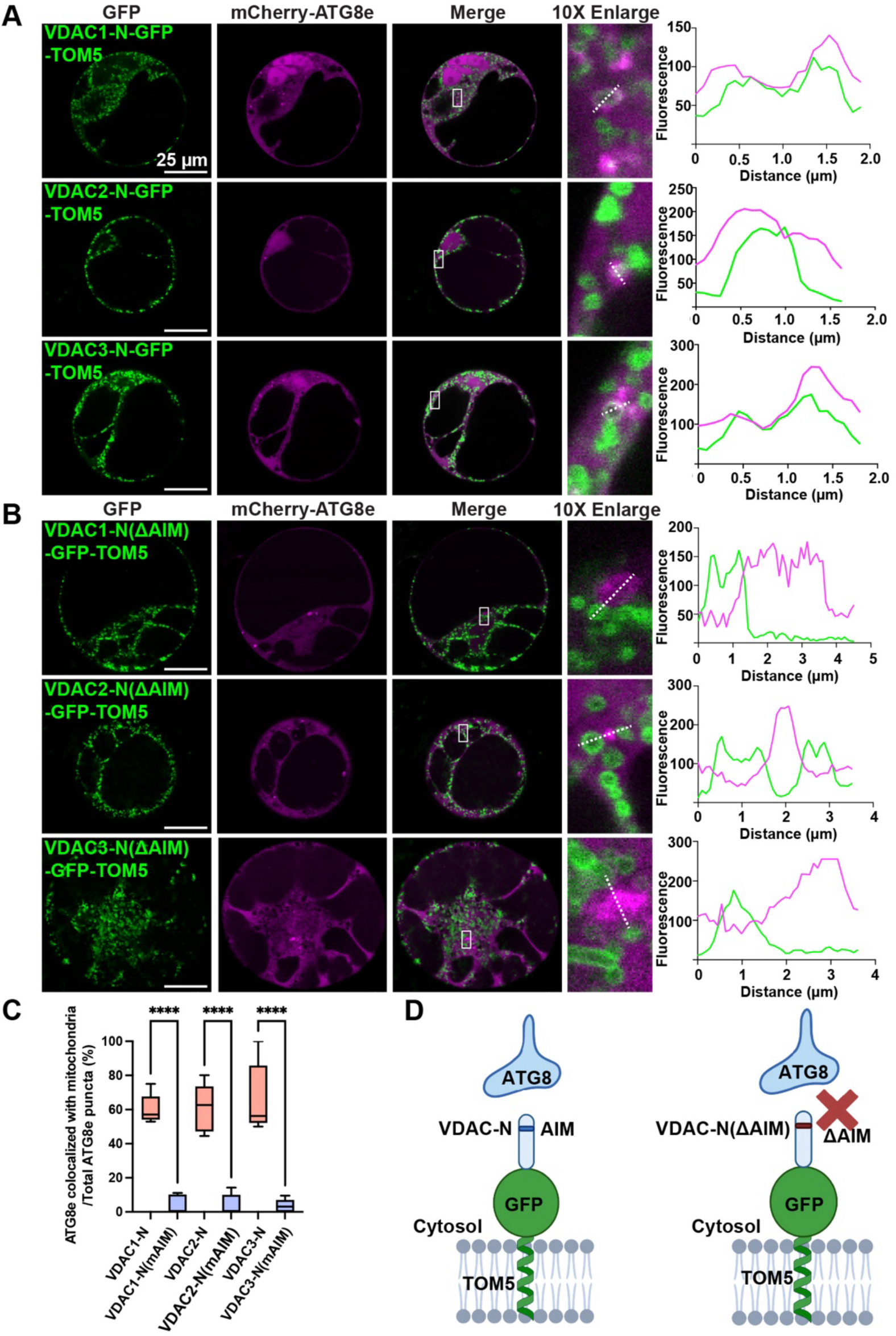
Artificial receptors containing the VDAC-N overhang recruit ATG8 to mitochondria in *Arabidopsis* protoplast. **A)** Confocal micrographs of *Arabidopsis* protoplasts co-expressing artificial outer mitochondrial proteins, VDAC-N-GFP-TOM5 and mCherry-ATG8e. The VDAC-N with the AIM is capable of recruiting mCherry-ATG8e to mitochondria when attached to GFP-TOM5. Scale bars: 25 µm. **B)** Confocal micrographs of *Arabidopsis* protoplasts co-expressing artificial outer mitochondrial proteins, VDAC-N-GFP-TOM5 without the AIM (VDAC-N(ΔAIM)-GFP-TOM5) and mCherry-ATG8e. The VDAC-N without AIM did not bring mCherry-ATG8e to mitochondria. Scale bars: 25 µm. **C)** Quantification of mCherry-ATG8e’s association with mitochondria when VDAC-N-GFP-TOM5 (A) or VDAC-N(ΔAIM)-GFP-TOM5 (B) were expressed. Unpaired t-test; *p* < 0.0001. **D)** Diagrams illustrating the structures of artificial receptors containing WT or ΔΑΙΜ versions of the VDAC N-terminus. GFP is attached to visualize mitochondria and TOM5 is employed for mitochondria targeting of the artificial receptors.

### Accumulation of aberrant mitochondria in *vdac* mutants

Mitochondria in T-DNA inserted mutant lines of VDAC1, 2, and 3 were compared with those in wild-type (Col-0) after DNP treatment. *Arabidopsis* seedlings expressing mitochondrial targeting marker Mito-GFP were pre-stained with TMRE to discern depolarized mitochondria that corresponded to green puncta due to the lack of magenta fluorescence from TMRE. *vdac* seedlings showed no difference from Col-0 in the control groups treated with DMSO. However, in experimental groups incubated with DNP, the percentages of depolarized mitochondria increased in the *vdac* seedlings (Figure 6A and 6C). Accumulation of depolarized mitochondria are more pronounced in *vdac* double and triple mutant lines than Col-0 (Figure 6C, Supplementary Figure S3, and Supplementary Figure S4). With more *VDACs* inactivated, the proportion of mitochondria without membrane potential increased in the absence of DNP, indicating that VDAC1, 2, and 3 are functionally redundant in the maintenance of mitochondrial electron transport chain. Moreover, we were able to rescue the defective membrane potential and accumulation of depolarized mitochondria in the *vdac* mutant lines by transforming them with normal copies of the *VDACs.* When AIM-mutated versions of VDACs were used for complementation experiments, they failed to revert the phenotype of depolarized metachronal accumulation in agreement with the role of the AIM in mitophagy (Supplementary Figure S6).

**Figure 6.**
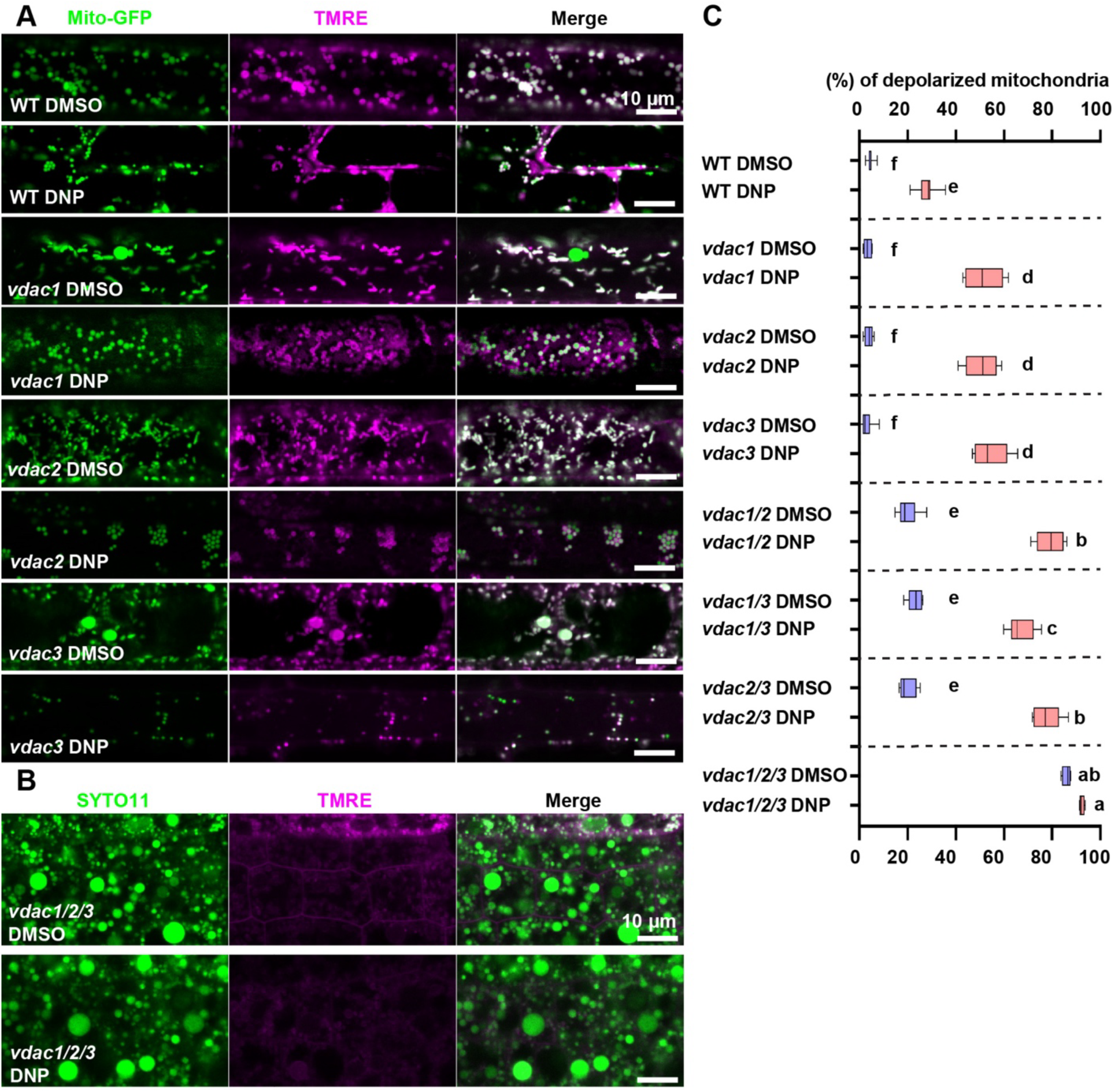
Depolarized mitochondria accumulated in *vdac* mutants when subjected to DNP-stress. **A)** DNP-induced mitochondria depolarization in wild-type (WT) and *vdac* seedling root cells. Mitochondria labeled with Mito-GFP were stained with a membrane potential sensitive dye, TMRE. White puncta correspond to normal mitochondria while depolarized mitochondria exhibit green fluorescence from Mito-GFP only. Scale bars: 10 µm. **B)** DNP-induced mitochondria depolarization in *vdac1/2/3* mutants. Mitochondria were stained with SYTO11 and TMRE. Normal mitochondria in the merged image panels are discerned by their white fluorescence while depolarized mitochondria exhibit only green fluorescence from SYTO11. Scale bars: 10 µm. **C)** Box-whisker plots illustrating percentages of depolarized mitochondria in WT, *vdac1*, *vdac2*, *vdac3-1*, *vdac1/2*, *vdac1/3*, *vdac2/3*, and *vdac1/2/3* root cells. Letters on the graph indicate significant difference (*p* < 0.05) calculated by one-way ANOVA followed by Tukey’s test.

Cristae collapse and electron-dense precipitates form in depolarized mitochondria when examined with electron microscopy (Zhou et al., 2016; Ma et al., 2021). Depolarized mitochondria accumulation was also evident in electron micrographs of Col-0 and *vdac* root cells after DNP treatment (Figure 7). Such mitochondria enclosed in phagophores (*i.e.* developing mitophagosomes) were noticed in electron micrographs of DNP-treated Col-0 cells. By contrast, mitophagosomes were rare in *vdac* cells and numerous depolarized mitochondria persisted in cytosol (Figure 7A-7B). To better visualize the difference between normal and damaged mitochondria, we carried out three-dimensional (3D) electron tomography (ET) imaging of mitochondria in *vdac1/2/3* triple mutants before and after DNP treatment in comparison to Col-0 mitochondria (Figure 7C). Diameters of Col-0 mitochondria were approximately 700 nm, with no electron density dark aggregates observed (Figure 7C & Movie 1). However, mitochondria in *vdac1/2/3* were swollen with diameters ranging from 1 - 2 µm and dark precipitates in the matrix (Figure 7C & Movie 2). DNP treatment worsened the mitochondria swelling in the triple mutant. Mitochondria with diameters longer than 3 µm were readily noticed. Their matrix often had large empty spaces, and their membrane integrity was compromised (Figure 7C & Movie 3). Altogether, absence of VDAC1, 2, and 3 delays the elimination of damage mitochondria, leading to the accumulation of depolarizing mitochondria. These findings suggest that the efficiency of *Arabidopsis* mitophagy is dependent on the three VDACs.

**Figure 7.**
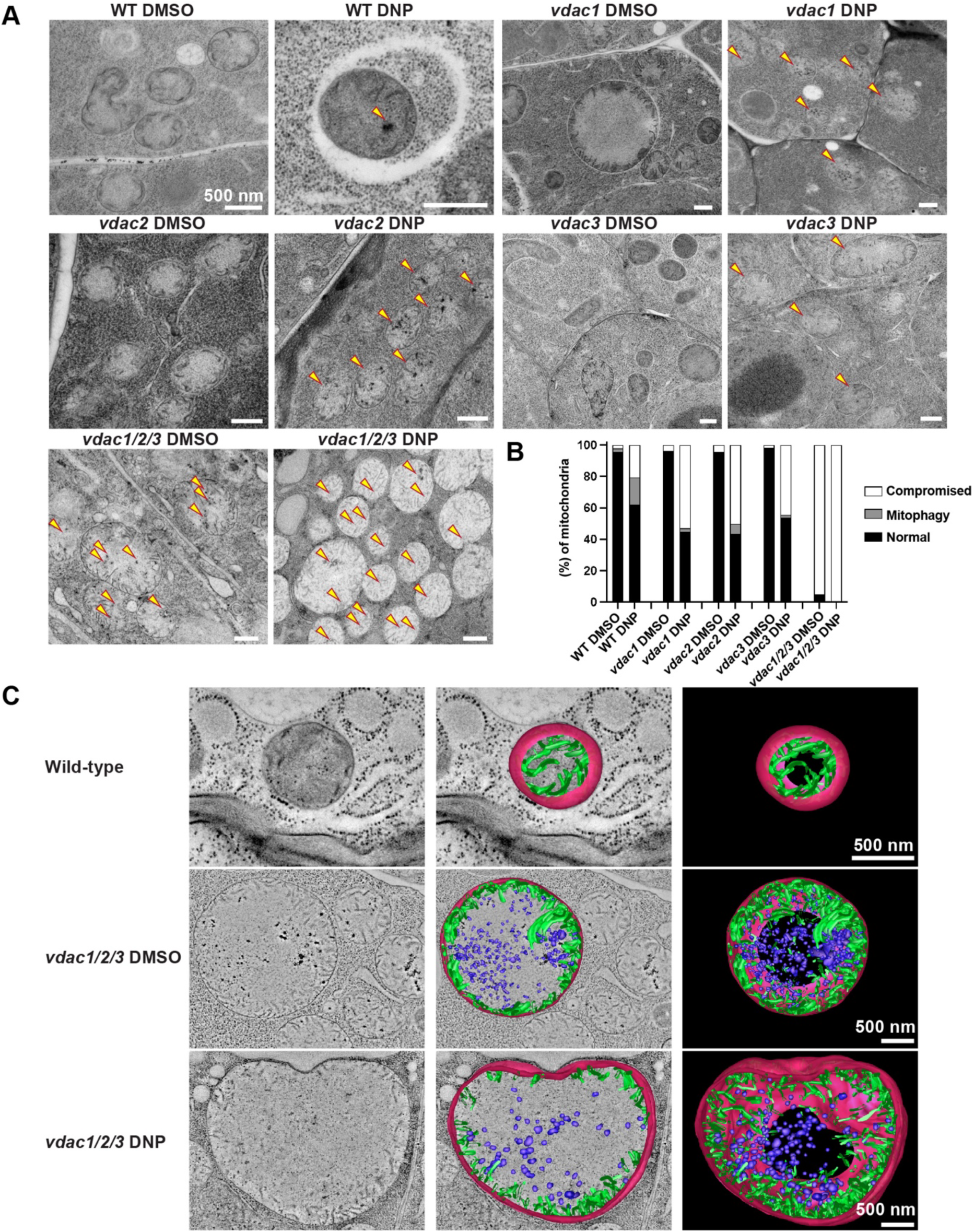
Accumulation of aberrant mitochondria in *vdac* mutants after DNP treatment. **A)** TEM micrographs of mitochondria in *Arabidopsis* WT, *vdac1*, *vadc2*, *vdac3-1* and *vdac1/2/3* seedling root cells treated with DMSO or DNP. Aberrant mitochondria with collapsed cristae distinctive of depolarized mitochondria are marked with yellow arrowheads. Scale bars: 500 nm. **B)** Bar charts illustrating percentages of normal mitochondria, damaged mitochondria, and mitophagosome-engulfed mitochondria in WT and *vdac1*, *vdac2*, *vdac3-1* and *vdac1/2/3* seedling root cells. **C)** 3D electron tomograms and models of mitochondria in WT, *vdac1/2/3* mutant, and DNP-treated *vdac1/2/3* mutant. Outer membrane, cristae connected to the inner membrane, and cristae aggregates were meshed into magenta, green, and blue surfaces. Scale bar: 500 nm.

### Mitophagosome formation is impaired in *vdac* mutants

Since aberrant mitochondria accumulated more in DNP-treated *vdac* cells than DNP-treated Col-0 cells (Figure 6 and 7), we examined whether mitophagy is affected by *VDAC* inactivation. *vdac* mutant lines co-expressing Mito-GFP and mCherry-ATG8e were generated to monitor mitochondria and autophagosomes. Ratios of mCherry-ATG8e puncta overlapping with mitochondria (i.e. mitophagosomes) to all mCherry-ATG8e puncta representing all types of autophagosomes were calculated to measure mitophagy activities (Figure 8A and 8B). In the absence of DNP, the three *vdac* mutant lines did not exhibit differences in the mitophagy activities from Col-0. Approximately 5-10 % of mCherry-ATG8e puncta were associated with mitochondria. However, DNP-treatment increased the ratio significantly in Col-0, but less increment in the three *vdac* lines. Among the *vdac* lines, mutation of *VDAC1* affected most as DNP treatment failed to increase the mitophagy activity (Figure 8B). When mitophagy flux was examined with immunoblot analyses, degradation of a mitochondrial matrix protein was delayed in the three *vdac* mutant lines (Figure 8C and 8D). Altogether, absence of VDAC will impact mitophagosome formation to remove depolarized mitochondria.

**Figure 8.**
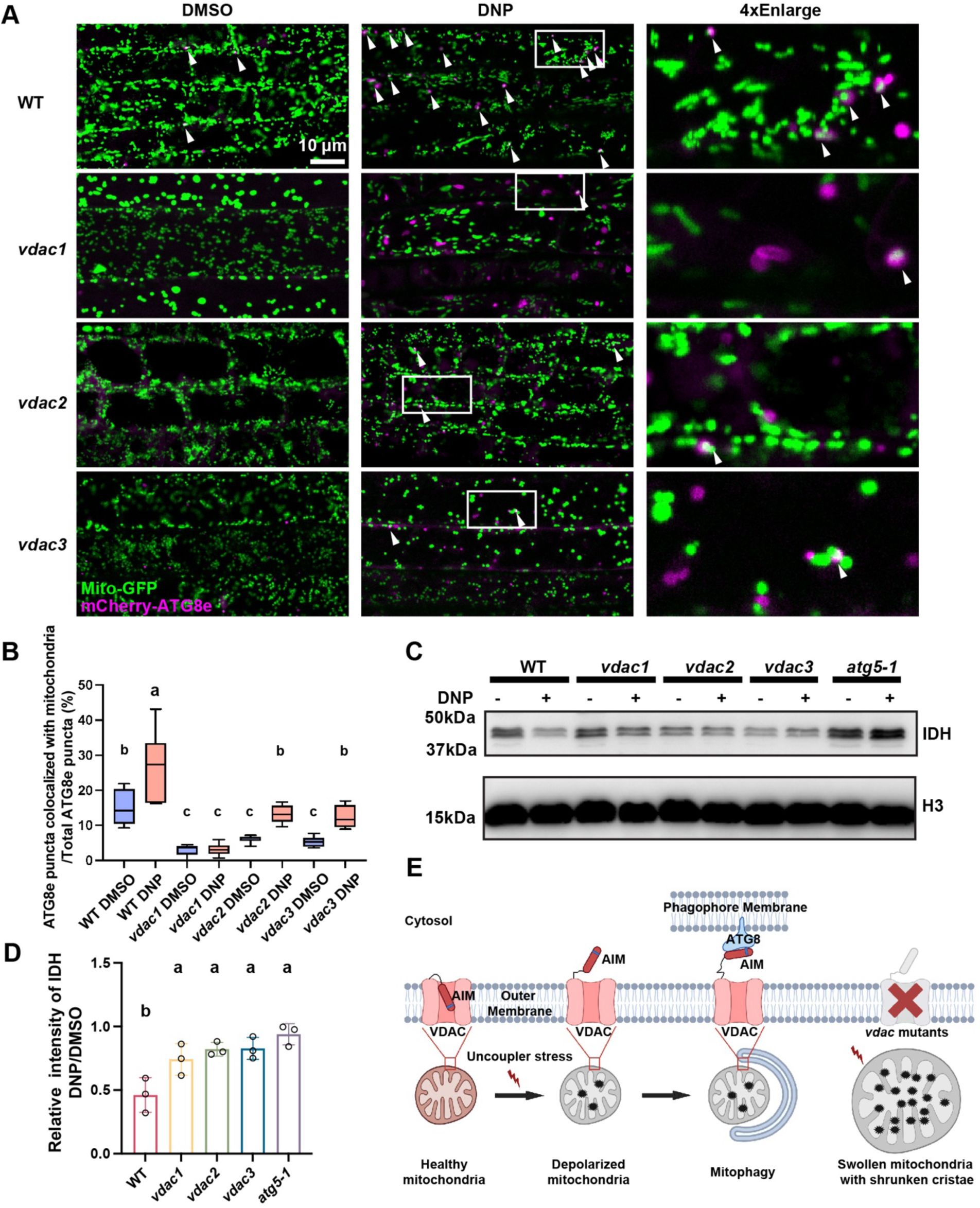
Mitophagy is impaired in *vdac* mutants. **(A)** Confocal micrographs of *Arabidopsis* WT, *vdac1*, *vdac2*, and *vdac3* seedling root cells expressing Mito-GFP and mCherry-ATG8e after DMSO- or DNP-treatment. Green puncta (Mito-GFP) associated with magenta spots (mCherry-ATG8e) were counted as mitochondria being targeted by autophagosomes. Such mitochondria are marked with white arrowheads. Scale bar: 10 µm. **(B)** Bar-whisker plots illustrating percentages of ATG8e puncta associated with mitochondria to all ATG8e puncta. More than eight seedling roots were examined for counting ATG8 spots under each condition. Three different cells per seedling root were imaged for counting. Letters in the graph denote significant difference (*p* < 0.05) calculated by one-way ANOVA followed by Tukey’s test. **C)** Immunoblot analyses of a mitochondrial matrix protein, IDH in *Arabidopsis* seedling root samples of wild-type, *vdac1*, *vdac2*, *vdac3*, and a*tg5-1* after DMSO- and DNP-treatment. Intensities of IDH were measured to compare mitochondria degradation induced by DNP. **D)** Quantification of the immunoblot analyses in **C)**. IDH amounts were normalized with amount of H3 for each genotype before calculating reduction in IDH levels caused by DNP (DNP/DMSO). Letters in the graph denote significant difference (*p* < 0.05) calculated by one-way ANOVA followed by Tukey’s test. n=3. **E)** VDAC-mediated mitophagy of depolarized mitochondria in *Arabidopsis*. Without VDACs (*i.e. vdac* mutant), aberrant mitochondria accumulate.

## Discussion

In this study, we identified three members of an *Arabidopsis* OMM protein family, VDAC1, 2, and 3 as mitophagy receptors. Notably, the N-terminal overhang of the each VDAC contains an AIM that facilitates the interaction with ATG8 when mitochondrial membrane potential is impaired (Figure 8E). While VDACs in mammals are ubiquitylated and function as substrates for mitophagy adaptors such as p62 (Geisler et al., 2010), our findings indicate that VDAC1, 2, and 3 of *Arabidopsis* directly bind ATG8 for the assembly of mitophagosomes (Figure 2 and 3). Conformational changes of VDAC N-terminal overhang from VDAC channel to cytosol in depolarized mitochondria is critical for ATG8 recruitment (Figure 4 and 5). Double or triple mutant lines of these VDACs accumulated more depolarized mitochondria and exhibited a delay in mitophagy compared to the single mutant lines (Figure 6, 7, and 8). This suggests that VDAC1, 2, and 3 have overlapping roles in the mitophagy process and sufficient VDACs are required by efficiency of mitophagy. These findings uncovered a plant-specific role of voltage-dependent anion channels of *Arabidopsis* in mitophagy, which has not been reported.

VDACs exist in open and closed states, with their gating mechanism closely linked to the mitochondrial membrane potential (Shuvo et al., 2016; Ngo et al., 2022). An increase in membrane potential typically results in the closure of the VDAC gate, while a reduction opens the channel. This gating behaviour is primarily governed by the N-terminal domain, which interacts with the β-barrel domains of VDACs, thereby determining whether the channel is in an open or closed state (Geula et al., 2012; Shuvo et al., 2016). There some research reported that VDAC N-terminus also undergo a translocation in stress condition, especially in apoptosis. They will translocate out of the channel and improve VDAC oligomerization and apoptosis (Shoshan-Barmatz et al., 2010; Kim et al., 2019). When isolated mitochondria were depolarized, we observed that the N-terminal overhang of *Arabidopsis* VDACs became approachable to proteinase K degradation, both *in vivo* and *in vitro* DMSO- or DNP-treatment are performed (Figure 4, and Supplementary Figure 2). This result suggests that the VDACs on mitochondria losing the membrane potential heightens accessibility of their N-terminal domains to cytosolic proteins. Given the VDACs have an AIM in their N-terminus, it is plausible that ATG8 is recruited to depolarized mitochondria via the exposed AIMs of VDACs, allowing for the selective targeting of those dysfunctional mitochondria for degradation.

In mammalian cells, VDACs on depolarized mitochondria undergo ubiquitylation, which promotes ATG8 recruitment to such mitochondria for degradation (Ham et al., 2020). Loss of mitochondria membrane potential activates PINK1 that phosphorylates Parkin which in turn add ubiquitin chains to its substrate, amplifying the alert signal, for promoting the recognition of damaged mitochondria (Harper et al., 2018). However, such signal amplifying pathway have not been uncovered in *Arabidopsis*. Furthermore, we could not find any evidence of additional VDAC ubiquitination induced by treatment with DNP (Supplementary Figure 7). The absence of depolarization-induced ubiquitination of VDACs suggests a divergent mechanism of mitophagy regulation in *Arabidopsis*.

VDACs are among the most abundant proteins constituting OMM, underscoring their significance in mitochondrial function and dynamics. VDACs account for more than 30% of OMM of *Arabidopsis* and VDAC1, 2, 3 are the three most abundant VDAC isotypes (Fuchs et al., 2020). We propose that VDAC1, 2, and 3 expose their AIMs to interact directly with ATG8 in the cytosol, priming depolarized mitochondria for the quick assembly of mitophagosomes once ATG8 becomes activated. This streamlined interaction could enable the efficient removal of damaged mitochondria without the need for extensive signal amplification. Further research will elucidate the mitophagy pathway in plants evolved for adapting to stresses distinct from mammalian cells. The interaction between AIMs and ATG8 proteins is influenced by both the core sequence of the AIM and the surrounding amino acid residues, highlighting the context-dependent nature of their interactions (Marshall et al., 2019; Wesch et al., 2020). *Arabidopsis* genome encodes five VDAC genes and VDAC4 and VDAC5 also contain an inferred AIM sequence, FADI, in their N-terminal domains. This AIM sequence is different from those of VDAC1, 2, or 3. Furthermore, VDAC4 uniquely possesses a proline residue in front of its AIM, while other VDACs feature a leucine at this position (Supplementary Figure 1). The deviation implies a distinct conformation surrounding the AIM of VDAC4. Although we failed to exclude the auto-activation of AD-VDAC3 in Y2H assay (Figure 2D), it still provides evidence that VDAC1, 2, and 3 interact with all the nine isotypes of *Arabidopsis* ATG8s. The copy number of VDAC5 is estimated to be only 0.07-0.13% of VDAC1-3, suggesting that its contribution to mitophagy is negligible (Fuchs et al., 2020).

In our study, we induced mitophagy using a depolarizing chemical rather than through natural stressors. Nevertheless, our data strongly support the involvement of VDAC1, 2, and 3 in the autophagic removal of depolarized mitochondria. Although these VDACs seem to play redundant roles in the DNP-induced mitophagy, it has been reported that they exhibit cell type-specific expression patterns, which may influence their contribution to the mitochondria quality control under varying environmental conditions (Robert et al., 2012). We also do not understand how the autophagy machinery upstream of ATG8 is activated following DNP treatment. Further research on how depolarized mitochondria communicate with upstream autophagic complex such as the ATG1 complex could offer clues.

## Acknowledgments

We appreciate Prof. Liwen Jiang (The Chinese University of Hong Kong) for pBI221-pUBQ10-GFP-GENE plasmid and PSBD cell. This work was supported by grants from the Research Grants Council of Hong Kong (GRF14113921, GRF14121019, GRF14109222, N_CUHK462/22, and C4002-20WF) to B-HK.

## Author Contributions

W.M., J.M., and B.-H.K. conceived and designed the experiments. W.M., and J.M., performed the IP-MS. W.M., J.M., K.Z., M.Z., and X.Zhuang. prepared the materials. X.Zheng. and J.Z. perform Y2H. WM took the confocal microscopy. J.M., W.M., and K.Z. isolated mitochondria and proceed proteinase K digestion. W.M. and P.W. carried out electron microscopy/tomography analysis. W.M. and K.Z., prepared 3D tomographic models. W.M. and J.M. did the immunoblot and pull-down experiments. W.M., K.Z., and L.F. performed transient expression. WM, JM, and B.-H.K. analyzed the data. W.M., and B.-H.K. wrote the manuscript.

## Methods

### Plant materials and growth and treatment conditions

The following plants have been previously described: YFP, YFP-ATG8e, Mito-GFP::mCherry-ATG8e, GFP-TOM5 (Kuhnert et al., 2020), *atg5-1* (SALK_145980) (Ma et al., 2021), *vdac1* (SALK_034653) (Pan et al., 2014), *vdac3-1* (SAIL_238_A01) (Kwon, 2016), and *vdac3-2* (SALK_127899) (Michaud et al., 2014). T-DNA insertional mutants *vdac2* (SAIL_342_D12) were obtained from the Arabidopsis Information Resource (TAIR) (www.arabidopsis.org). pCAMBIA1300-p35S::GFP-VDAC (WT/ΔAIM) were transfected into *vdac* mutants via Agrobacterium to construct complementary lines. Maps and genotyping of *vdac* mutants are present in Supplementary Figure 5. Double mutant *vdac1/vdac2*, *vdac1*/*vdac3*, and *vdac2*/*vdac3* are crossed from *vdac1* and *vdac2*, *vdac1* and *vdac3-1*, *vdac2* and *vdac3-2*, respectively. Triple mutant *vdac1/2/3* are achieved via crossing from *vdac1*, *vdac2* and *vdac3-2*. Mito-GFP/mCherry-ATG8e/*vdacs* are achieved via crossing between Mito-GFP/mCherry-ATG8e/Col-0 with *vdac* single mutants. For plant growing, seeds were sterilized with 70% ethanol and then plated on medium containing half-strength Murashige and Skoog (MS) salts and 0.8% agar. Seedlings were grown at 22 °C under a 16-hour light/8-hour dark photoperiod. For autophagy induction, 7-day-old seedlings were transferred to liquid half-strength MS medium containing DMSO (1:200) as a control or 50 μM DNP for 2 hours. After DNP treatment, the seedlings were washed with half-strength MS medium and incubated for 0.5 to 1.5 hours before observation or as indicated.

### Plasmid construction

For the plasmids used in this study, complementary DNAs (cDNAs) from *Arabidopsis* seedlings encoding the target genes were amplified using high-fidelity PCR and cloned into the corresponding plasmids either through restriction digestion (New England BioLabs) or recombination cloning (Vazyme, ClonExpress Ultra One Step Cloning Kit V2, C116-02). For transgenic plants, full-length cDNAs were amplified and inserted into the pBI121-p35S::GFP-Gene plasmid via restriction digestion using BamHI, KpnI, and T4 ligase. For transient expression, the wild-type or AIM-mutated version of VDAC was cloned into the modified pBI221-pUBQ10::GFP-Gene vector using SpeI, XhoI, and T4 ligase. For yeast two-hybrid (Y2H) and pull-down assays, the wild-type or site-mutated versions of VDAC and ATG8 were inserted into pGADT7, pGBKT7, pGEX-KG, and pET30a(+) vectors using restriction digestion. For artificial receptor construction, VDAC-N or VDAC-N(ΔAIM), along with GFP and TOM5, were cloned into the pBI221-pUBQ10 backbone using a one-step recombination cloning kit.

### *In vivo* co-immunoprecipitation

Protein extraction and immunoprecipitation were performed as described previously (Ma et al., 2021). 7-day-old *Arabidopsis* seedlings were incubated in 50 µM DNP for 2 hours, followed by a 1-hour washout period before protein extraction. Total plant lysates were centrifuged at 14,000 rpm for 10 minutes at 4°C. The supernatant was prepared in lysis buffer (50 mM Tris/HCl, pH 7.4; 150 mM NaCl; 0.5 mM EDTA; 5% glycerol; 0.2% CA-630; 2 mM DSP (dithiobis(succinimidyl propionate)); and 1x Complete Protease Inhibitor Cocktail) and incubated with BeyoMag™ Anti-His Magnetic Beads (Beyotime, P2135) for 4 hours at 4°C. The beads were then washed five times with wash buffer (25 mM Tris/HCl, pH 7.4; 150 mM NaCl; and 0.5 mM EDTA, with 1x Complete Protease Inhibitor Cocktail) and eluted by boiling in 2× Laemmli buffer. The samples were separated by SDS-PAGE and analyzed by immunoblotting using the indicated antibodies.

### IP-MS and Gene Ontology analysis

The Co-IP process for HPLC-mass spectrometry analysis was performed as described above. Lysates from free YFP and YFP-ATG8e were pulled down using Anti-GFP beads (ChromoTek, gtma-20). The protein samples were boiled with 2× Laemmli buffer, slightly separated on an SDS-PAGE gel, and then stained with Imperial™ Protein Stain (ThermoFisher, 24615). In-gel digestion was performed as described (Zhou et al., 2023).

Trypsin-digested protein samples were analyzed using Orbitrap Fusion Lumos Tribrid mass spectrometer (ThermoFisher). The raw data was processed with MaxQuant 2.4 and Perseus software (https://www.maxquant.org/), using TAIR10 database (https://www.arabidopsis.org/) for protein identification (Tyanova et al., 2016). Volcano plot, Venn diagram, and Pie charts were generated using Microsoft Excel (https://www.microsoft.com/) and Prism 10 (https://www.graphpad.com). Gene Ontology analysis for cellular component was conducted using the Gene Ontology platform (http://geneontology.org/) (Ashburner et al., 2000).

### Confocal microscopy and image processing

Confocal fluorescence images were acquired using a Leica Stellaris 8 FALCON confocal system with a 63× water-immersion lens (Leica Microsystems). 7-day-old *Arabidopsis* seedlings were incubated in DMSO (1:200) or 50 µM DNP in liquid half MS medium for 2 hours and wash out drugs for 1 hour before imaging. 1 µM Tetramethyl rhodamine ethyl ester (TMRE, ThermoFisher, T669) and 2.5 µM SYTO11 (ThermoFisher, S7573) were applied to stain *Arabidopsis* root cell mitochondria for 10 mins. Sequential acquisition was used when observing fluorescent markers. Images were processed with Adobe Photoshop software (https://www.adobe.com) and the microscopy data were evaluated with one-way ANOVA tests using Prism 10 (https://www.graphpad.com).

### Yeast two-hybrid assay

The full-length CDS of ATG8a-i were cloned into pGBKT7 vector to fuse with the GAL4 DNA binding domain. The full-length CDS of VDAC1, VDAC1(ΔAIM), VDAC2, VDAC2(ΔAIM), VDAC3, VDAC3(ΔAIM) were inserted into pGADT7 vector to fuse with the activating domain.

### *In vitro* pull-down assays

Recombinant proteins were expressed using the BL21(DE3) *E. coli* strain. Bacteria were grown to an OD_600_ of 0.6–0.7, followed by induction with 500 μM IPTG and incubation for over 20 hours at 16°C. 10 mL of culture were pelleted and resuspended in 2 ml of EDTA-free buffer (25 mM Tris/HCl, pH 7.4; 150 mM NaCl; 5% glycerol; 1 tablet/50 ml cOmplete, EDTA-free Protease Inhibitor Cocktail [Sigma-Aldrich]; with detergent added when necessary). Cells were lysed through sonication (Ultrasonic Processor 130 W [Sigma-Aldrich], Z412619; 2 minutes, 2-second pulse on, 2-second pulse off, 75% amplitude). Lysates were clarified by centrifugation at 15,000 rpm at 4°C for 10 minutes. The *E. coli* lysate was then mixed with 10 μl of equilibrated Anti-GST Magnetic Beads (Beyotime, P2138) and incubated for 2 hours at 4°C on a spinning wheel. The beads were washed five times with EDTA-free buffer, then eluted in 50 μl of 2× Laemmli buffer, and boiled for 10 minutes at 95°C. SDS-PAGE gels were loaded with 5 μl of sample per well. GST rabbit monoclonal antibody (Beyotime, AF2299) and Anti-ATG8a-i antibody (Agrisera, AS14 2811) were used at a 1:4000 dilution.

### Transient expression in *Arabidopsis* protoplasts

Transient expression in *Arabidopsis* PSBD cultures was performed as described previously (Miao and Jiang, 2007; Li et al., 2021). In brief, pBI221-pUBQ10 plasmids containing the gene of interest are prepared using the NucleoBond® Xtra Maxi Kit (MACHEREY-NAGEL, 740414.50). These plasmids are then introduced into PSBD cells via electroporation. The transfected protoplasts are incubated for 12 to 16 hours before being subjected to confocal imaging.

### AlphaFold2-multimer simulation

AlphaFold2-multimer protein interaction simulation platform was reported before.(Ibrahim et al., 2023) AF2-multimer was used through a subscription to the Google Colab (https://colab.research.google.com/github/sokrypton/ColabFold/blob/main/AlphaFold2.ipynb#scrollTo=svaADwocVdwl). Amino acid sequences of VDAC and ATG8 (wild-type and mutated version) were uploaded to the website. Predicted protein structure are present by ChimeraX-1.5 (https://www.cgl.ucsf.edu/chimerax/).

### Mitochondria isolation*, in vitro* or *in vivo* uncouple treatment and proteinase K digestion

Mitochondria were isolated via discontinuous density gradients as reported (Monika W. Murcha, 2015). Percoll solutions were prepared with percoll and 2x wash buffer (0.6 M mannitol, 20 mM TES N-Tris(hydroxymethyl)methyl-2-aminoethanesulfonic acid, 0.2 % (w/v) BSA, pH 7.5). Gradients were prepared by layering 1ml of 21 % (v/v) solution over 250 μl of 40 % (v/v) solution, continuing with 500 μl of 16 % (v/v) solution on the top. 7-day-old GFP-TOM5 *Arabidopsis* seedlings were ground with pestle in grinding buffer (0.3 M mannitol, 50 mM tetrasodium pyrophosphate, 2 mM EDTA, 0.5 % (w/v) PVP-40, 0.5 % (w/v) BSA, pH 8.0) for mitochondrial extraction. Lysates were decanted into 2ml tubes and centrifuged at 3500 × g at 4 °C for 5 minutes. The supernatant was transferred into a new tube and centrifuged at 18000 × g at 4 °C for 15 minutes. The pellet was resuspended gently in 1 ml 1x wash buffer and then centrifuged at 1000 × g at 4 °C for 5 minutes. The supernatant was transferred into a new tube and centrifuged at 15000 × g at 4 °C for 15 minutes. The pellet was resuspended thoroughly in 200 μl 1x wash buffer and then layered over Percoll gradients following centrifuging at 40,000 × g for 45 min without brake. The isolated mitochondria were in the faint yellow layer between 21 % (v/v) and 40 % (v/v) Percoll gradient solution. Finally, isolated mitochondria were carefully aspirated into a new tube for confocal microscopy imaging and proteinase K digestion assay. Purified mitochondria were separated into serval tubes, 50 µl each. DMSO and DNP were added with the same concentration to treatment on seedlings. Proteinase K solution (Invitrogen, No.1622076) was added to mitochondria with indicated concentrations. Digestions were processed on ice for 30 min. 1% Triton X-100 were used to fragment mitochondria membrane.

### TEM analysis, electron tomography, and 3D modeling

For TEM sample preparation, high-pressure freezing, freeze substitution, resin embedding, and ultramicrotomy were performed as previously described (Byung-Ho, 2010). Briefly, 7-day-old *Arabidopsis* seedlings were incubated in DMSO or DNP for the specified durations and then rapidly frozen using an HPF ICE high-pressure freezer (Leica Microsystems). The samples were freeze-substituted at -80°C for 72 hours, with excess OsO4 removed by rinsing with precooled acetone. After gradually warming up to room temperature over 48 hours, root samples were separated from the planchettes and embedded in Embed-812 resin (Electron Microscopy Sciences, Cat. No. 14120). Thin sections (100 nm) from each time point were prepared and examined using a Hitachi 7400 TEM (Hitachi High-Technologies) operated at 80 kV.

For dual-axis tomography analysis, semi-thick sections (250 nm) were collected on formvar-coated copper slot grids (Electron Microscopy Sciences) and stained with 2% uranyl acetate in 70% methanol followed by Reynolds’ lead citrate. Tilt series were acquired from 60° to -60° in 1.5° increments using a 200-kV Thermo Scientific™ Talos™ Arctica™ Cryo-TEM. Tomograms were reconstructed following the method described by Toyooka and Kang (Toyooka and Kang, 2014). Model generation utilized the auto-contour command (https://bio3d.colorado.edu/imod/doc/3dmodHelp/autox.html) in the 3dmod software package.

### Protein extraction and immunoblot analysis

Isolation of mitochondrial proteins were carried out as described previously (Ma et al., 2021). Briefly, 7-day-old *Arabidopsis* seedlings were incubated for 2 hours in DMSO (Sigma-Aldrich, D8418) or 50 µM DNP (Sigma-Aldrich, D198501) in liquid half MS medium (Caisson, MSP01) and then washed. Protein samples were analyzed by 15% SDS-PAGE. Primary and secondary antibodies were diluted in 1x phosphate-buffered saline (PBS). Antibodies against GFP (Abcam; ab290), mCherry (Abcam; ab167453), ATG8 (Agrisera; AS124769), isocitrate dehydrogenase (IDH; Agrisera; AS06203A), and H3 (Agrisera; AS10710) were used as indicated. Band intensities from three replicate immunoblot analyses were quantified using ImageJ (https://imagej.nih.gov/ij/) and Microsoft Excel (https://www.microsoft.com/). Graphs were prepared with Prism 10 (https://www.graphpad.com). Representative results from at least three independent immunoblot analyses are shown in the figures.

### Quantification and Statistical Analysis

All statistical comparisons were based on data from three or more independent biological replicates (indicated in figure legends), with similar variances observed between groups. Statistical analyses and graphing were using Prism 10 (https://www.graphpad.com). For data representing a single time-point and condition, we performed statistical comparisons between different groups using one-way ANOVA with Tukey’s multiple comparisons test. Two-tailed unpaired Student’s t test was performed to compare the difference between two groups. \**p* < 0.05, \*\**p* < 0.01, \*\*\**p* < 0.001, \*\*\*\**p* < 0.0001, n.s., no significant difference. Letters on the graph indicate significant difference (p < 0.05).

---

**Supplementary Figure 1.**
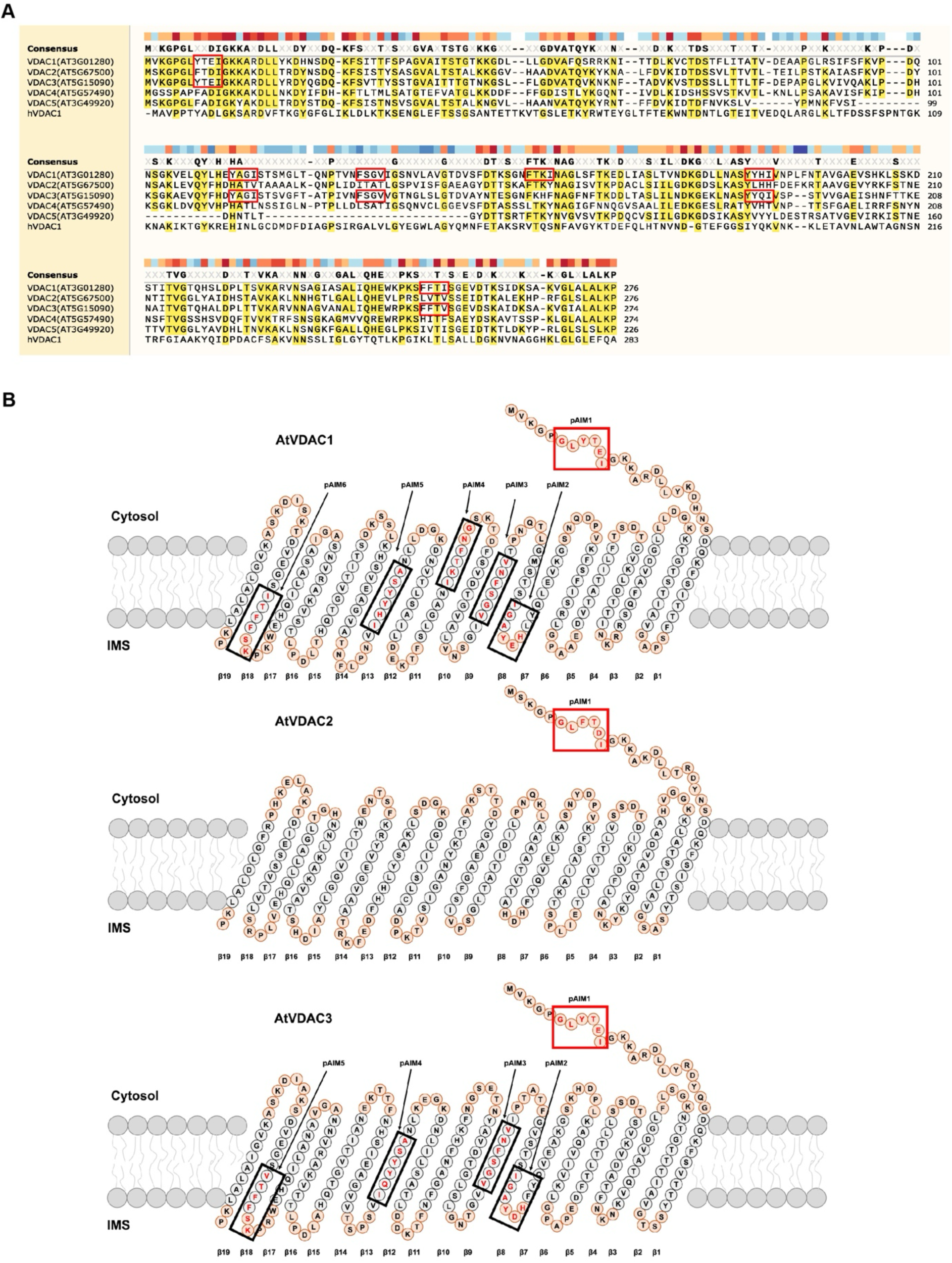
AIM prediction based on VDAC amino acid sequence analysis. **A)** Amino acid sequence alignment by the Snapgene software. All potential AIMs (pAIMs) in VDACs predicted by the iLIR website are highlighted with red boxes. **B)** Predicted topologic structures of AtVDAC1, AtVDAC2, AtVDAC3. All potential AIMs in VDACs in a are marked with box and amino acids in the AIMs are highlighted (red letters). AIMs in the N-terminal overhangs are highlighted with red box. IMS: mitochondrial inter membrane space.

**Supplementary Figure 2.**
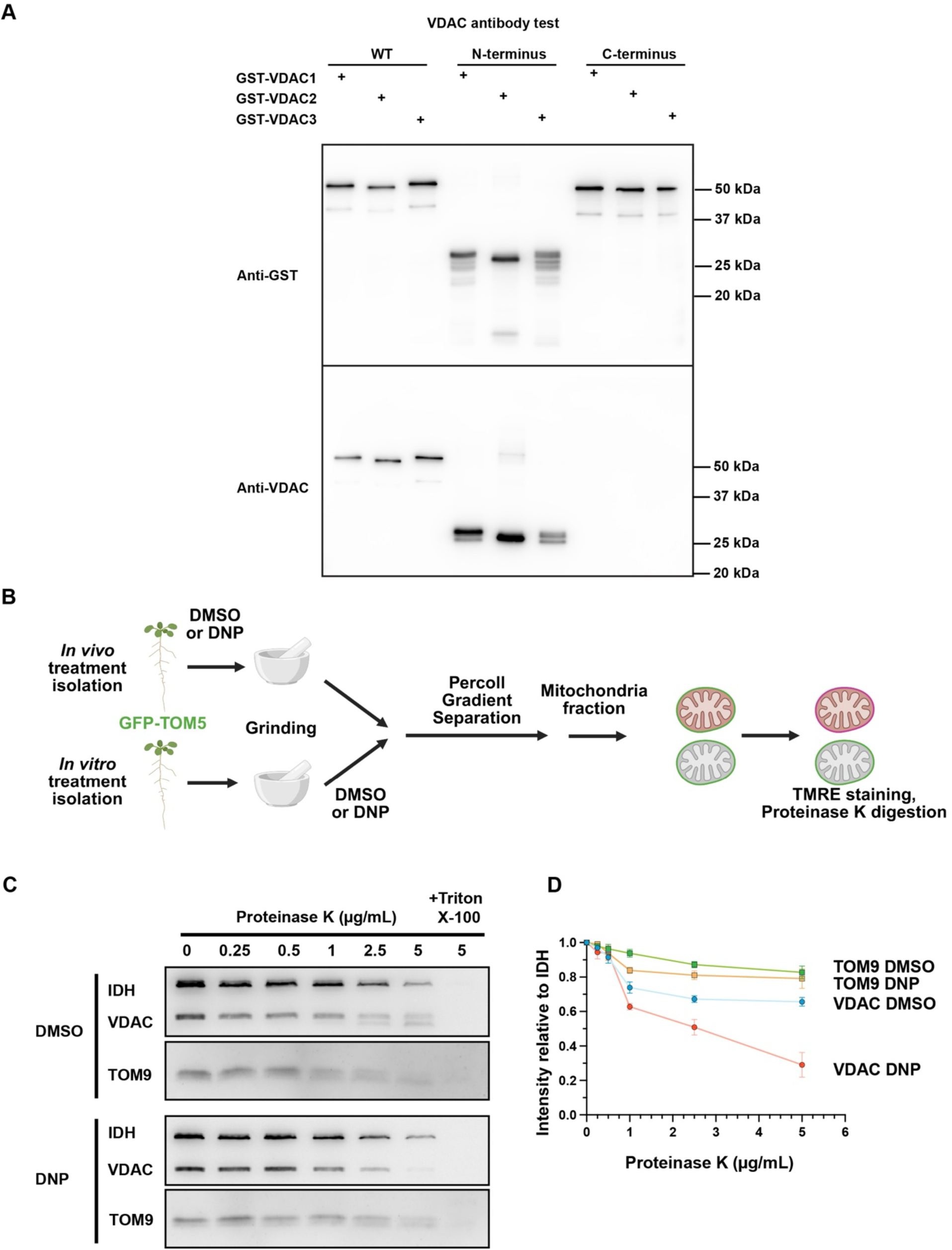
Characterization of the VDAC antibody for proteinase K assay. **A)** GST-fused VDACs, including full lengths, N-terminus (1-26 aa), C-terminus (27-C terminus) were purified from *E. coli.* lysates and incubated with Anti-GST (Beyotime, AF2299), and Anti-VDAC1-5 (Agrisera, AS07212). All purified proteins were detected by GST antibody. However, VDAC antibody recognized WT and N-terminus versions of GST-VDAC but not the GST-C-terminus version, indicating that the antibody is specific to the N-terminal 1-26 aa peptide. **B)** Diagram of *in vivo* or *in vitro* DMSO-/DNP-treatment *Arabidopsis* mitochondria isolation. **C)** Immunoblot analyses mitochondrial proteins after proteinase K digestion. Isolated mitochondria were treated with DMSO and DNP for 2 hours and digested by varying concertation of proteinase K. 1% Triton X-100 are added to dissolve mitochondrial membrane. **D)** Quantification of immunoblot in **C)**. Intensities of IDH were adopted for estimating amounts of mitochondria for each experimental sample. TOM9 amounts indicate the integrity of mitochondria outer membrane. All protein amounts relative to IDH were normalized to the results of the 0 µg/ml Protease K sample. Means ± s.d.. n=3.

**Supplementary Figure 3.**
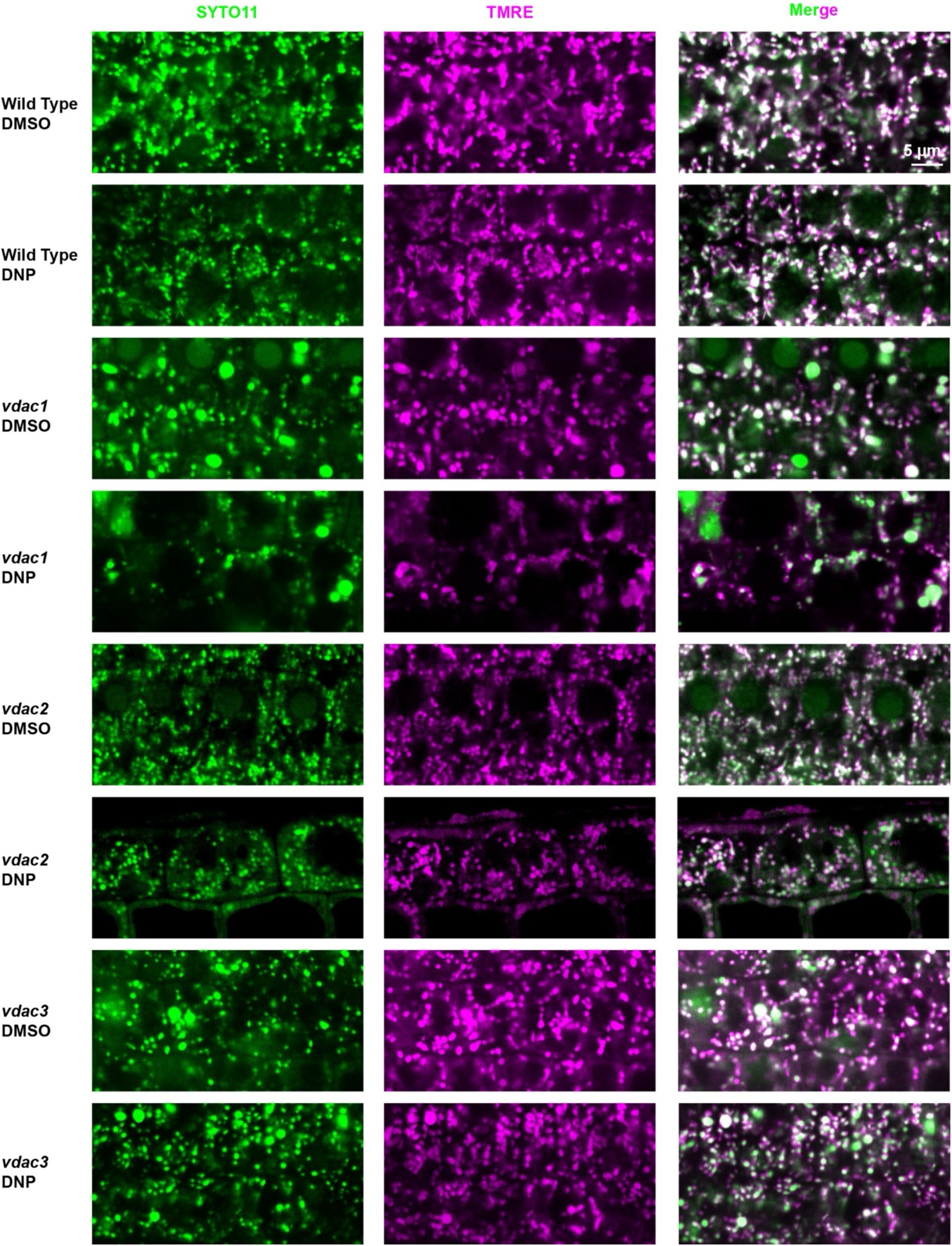
Mitochondria stained with SYTO11 and TMRE in WT and *vdac* single mutants with or without DNP stress. Mitochondria are visualized by SYTO11 that stains mitochondrial DNA. Their membrane potential levels were estimated with TMRE. Normal mitochondria are seen as white puncta from overlapping fluoresced from SYTO11 and TMER. Green puncta lacking TMRE fluorescence correspond to depolarized mitochondria. Scale bar: 5 µm

**Supplementary Figure 4.**
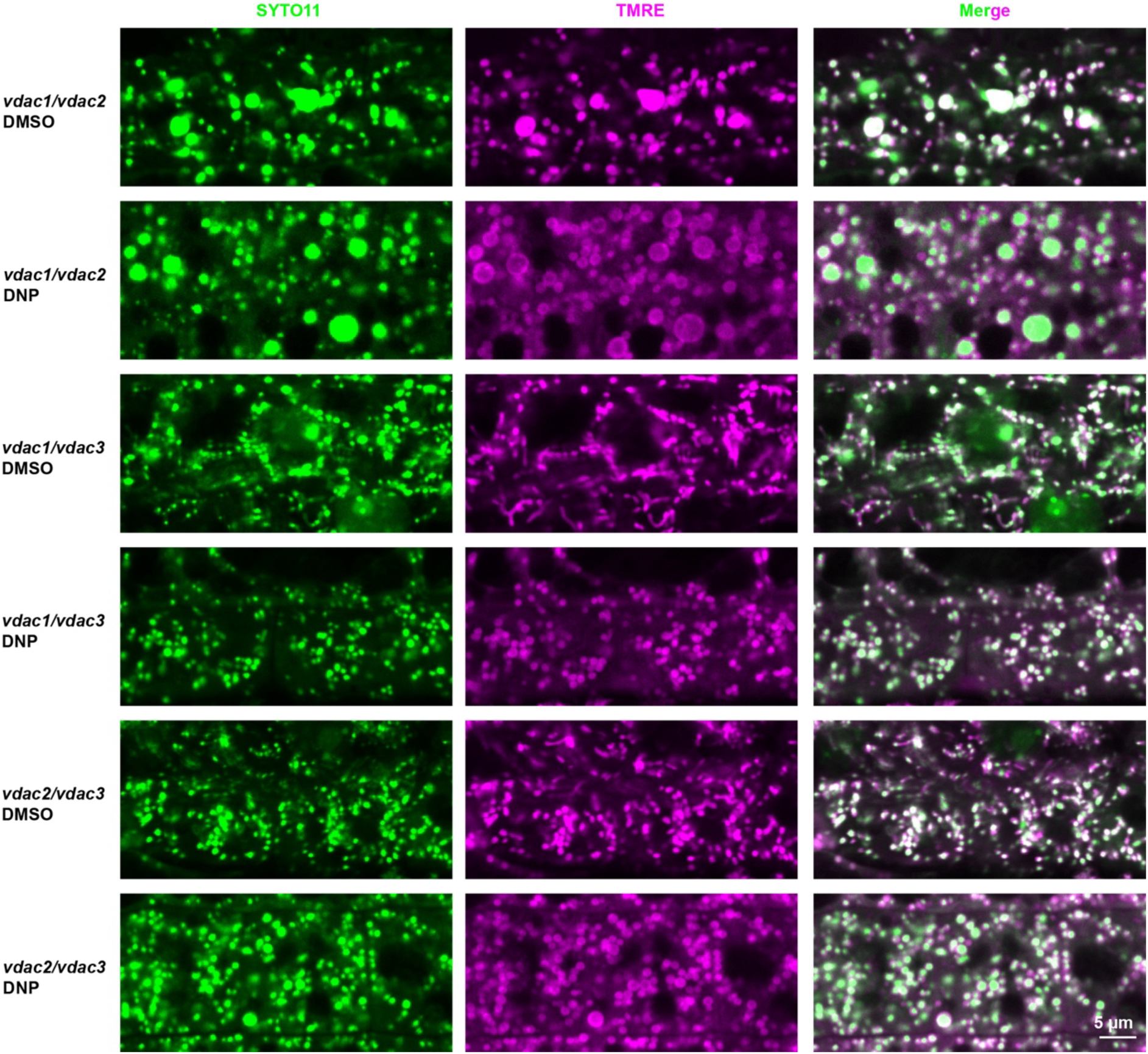
Mitochondria stained with SYTO11 and TMRE in WT and *vdac* double mutants with or without DNP stress. Mitochondria are visualized by SYTO11 that stains mitochondrial DNA. Their membrane potential levels were estimated with TMRE. Normal mitochondria are seen as white puncta from overlapping fluoresced from SYTO11 and TMER. Green puncta lacking TMRE fluorescence correspond to depolarized mitochondria. Scale bar: 5 µm.

**Supplementary Figure 5.**
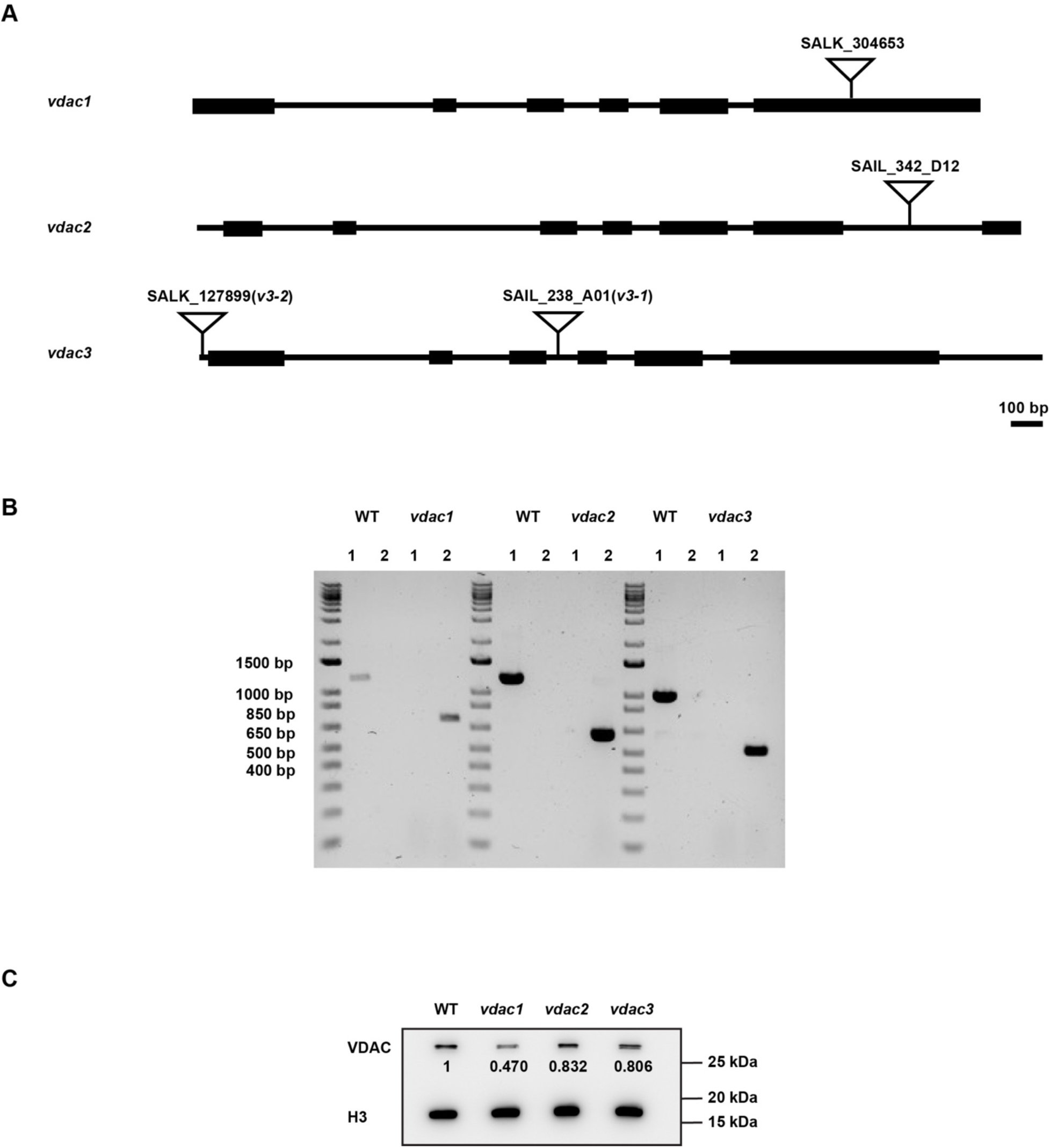
Characterization of *vdac* KO line. **A)** Schematic representation of T-DNA insertions in *vdac1, 2,* and *3*. **B)** PCR genotyping of *vdac1, vdac2,* and *vdac3-1* to verify T-DNA insertions. Lane 1: LP+RP; Lane 2: BP+RP. **C)** VDAC amount in WT, *vdac1*, *2*, and *3*. All VDAC level were normalized with those of H3 in same genotype, then all relative intensity were compared to that in WT.

**Supplementary Figure 6.**
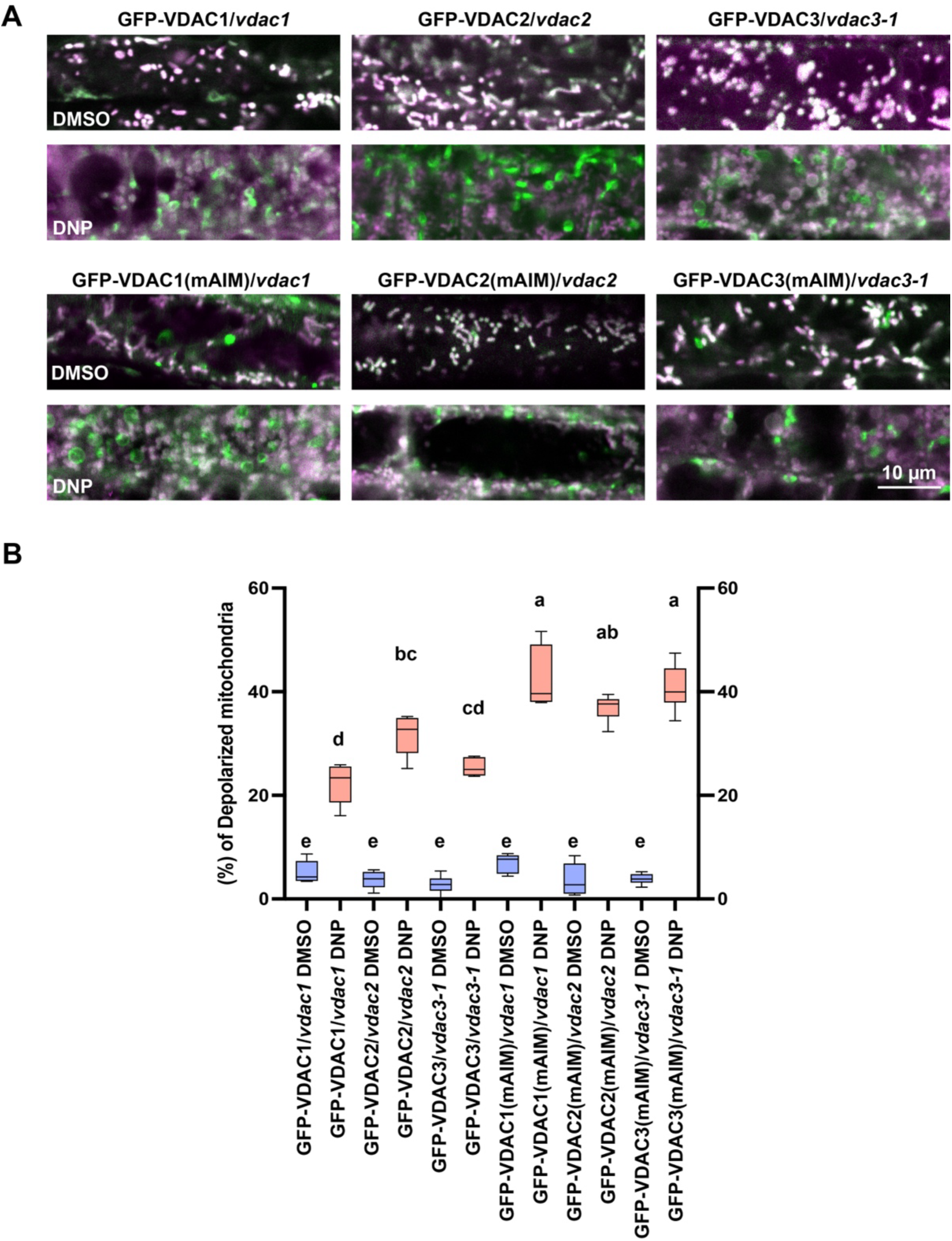
Complementation of *vdac1, 2, and 3*. **A)** GFP-fused VDAC1, 2, or 3 (WT/mAIM) were expressed in the respective *vdac* mutant lines and the transgenic lines were stained with membrane potential dye TMRE. Normal mitochondria are seen as white puncta from overlapping fluorescence from SYTO11 and TMRE. Green puncta lacking TMRE fluorescence correspond to depolarized mitochondria. p35S::GFP-VDACs (WT/mAIM) rescued the phenotypes of swelling mitochondria in *vdac1* and *vdac3*. Note that enlarged mitochondria disappeared in the mutant lines. Scale bar: 10 µm. **B)** Box-whisker plot illustrating percentages of depolarized mitochondria in GFP-VDACs (WT/mAIM) complemented *vdac1*, *vdac2*, *vdac3-1*, root cells. Letters on the graph indicate significant difference (*p* < 0.05) calculated by one-way ANOVA followed by Tukey’s test.

**Supplementary Figure 7.**
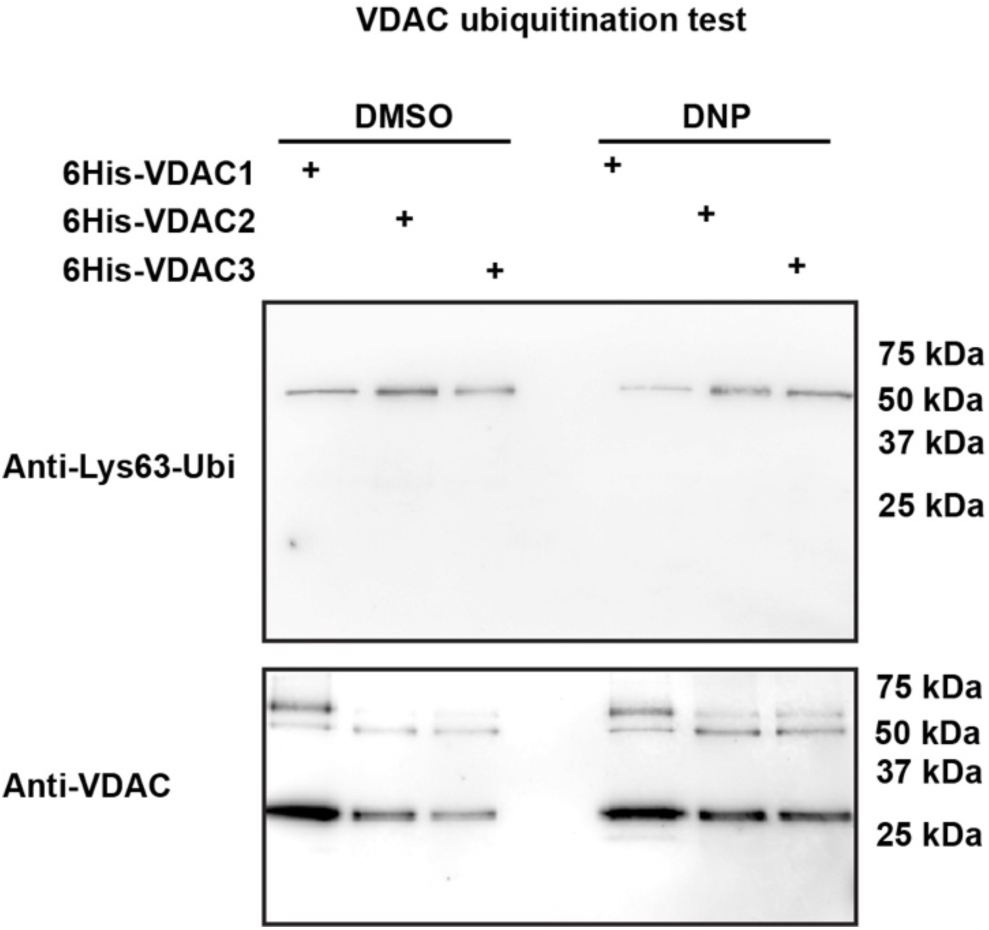
No changes in the ubiquitylation of VDACs with or without DNP treatment. 6×His-VDAC1, VDAC2, and VDAC3 are purified from *Arabidopsis* lysates using Ni-NTA beads. VDAC proteins were detected by the anti-VDAC antibody and ubiquitylations are checked by Anti-Ubiquitin Lys63-specific antibody (Sigma-Aldrich, clone Apu3, No.05-1308). Increases in sizes expected from ubiquitylation were not observed in VDACs after DNP treatment (anti-VDAC panel). No alterations in Lys63 ubiquitylation were not noticed either after DNP treatment (Anti-Lys63-Ubi).

**Movie 1. Electron tomogram of mitochondrion in DMSO-treated WT.**

Mitochondrial outer membrane and inner membrane were outlined by magenta and green, respectively. Scale bar: 500 nm.

**Movie 2. Electron tomogram of mitochondrion in DMSO-treated *vdac1/2/3*.**

Mitochondrial outer membrane, inner membrane, and aggregation of cristae were outlined by magenta, green, and blue, respectively. Scale bar: 500 nm.

**Movie 3. Electron tomogram of mitochondrion in DNP-treated *vdac1/2/3*.**

Mitochondrial outer membrane, inner membrane, and aggregation of cristae were outlined by magenta, green, and blue, respectively. Scale bar: 500 nm.

